# Interaction of IRS2 with PLK1 protects cells from mitotic stress

**DOI:** 10.1101/2025.07.17.665461

**Authors:** Ji-Sun Lee, Quyen Thu Bui, Minjeong Jo, Jennifer S. Morgan, Isha Nasa, Arminja N. Kettenbach, Leslie M. Shaw

## Abstract

In this study we identify a role for IRS2 in the protection of cells from mitotic stress through its interaction with PLK1. IRS2 is an adaptor protein for the insulin and IGF-1 receptors that mediates their signaling functions. In this capacity, IRS2 is tyrosine phosphorylated to recruit signaling effectors that control cellular outcomes. A role for IRS2 in mitotic regulation has been reported, but the mechanism of IRS2 action in this regulation has not been determined. Here we report that IRS2 interacts with PLK1 in a CDK1-dependent manner, and they co-localize at centrosomes in mitotic cells. In response to mitotic stress, cells that lack IRS2 or express a PLK1-binding deficient mutant exhibit reduced centrosome separation and a shortened mitotic arrest that leads to reduced tumor cell viability. In contrast, cells expressing an IRS2 mutant that is not tyrosine phosphorylated display normal mitotic function. Together, our findings establish a mechanistic connection between IRS2 and mitotic regulation that is distinct from its function as a signaling adaptor protein.

## Introduction

Insulin receptor substrate 2 (IRS2) is a member of the IRS family of adaptor proteins that mediate downstream signaling from the insulin (IR) and insulin-like growth factor (IGF1R) receptors in response to engagement of these receptors by their ligands (insulin, IGF1 and IGF2)^1^. Upon recruitment to the activated IR or IGF1R, the IRS proteins are phosphorylated on tyrosine residues by the intrinsic receptor kinases, generating binding sites for effectors to activate downstream signaling and mediate cellular outcomes ^2^. The IRS proteins have been shown to play essential roles in the regulation of normal metabolic homeostasis and cell growth by the insulin and IGF signaling (IIS) pathway, with each IRS protein mediating overlapping and distinct functional outputs ^3, 4^. In cancer, the IRS proteins have also been implicated in divergent functions. IRS1 and IRS2 both support IIS-dependent glucose uptake and cell survival and this can be dependent upon tissue-specific expression of each adaptor protein ^2^. However, proliferation is induced by IIS only when cells express predominantly IRS1, whereas cells expressing predominantly IRS2 migrate in response to the same signaling input ^5, 6^. IRS2 also regulates breast cancer stem cell function in response to IIS through the PI3K-dependent regulation of MYC stability ^7^. Recently, IRS2 was identified as a substrate of the anaphase-promoting complex/cyclosome (APC/C)^CDH1^ ubiquitin ligase complex that is activated late in mitosis to allow completion of cell division and return to the G_1_ cell cycle phase ^8^, revealing a new role for IRS2 in mitosis.

Cell cycle progression is a complex and tightly regulated process which is coordinated by the activation of multiple kinases, regulation of gene expression and targeted protein degradation^9^. Progression is monitored by checkpoints within each cell cycle stage to ensure cellular fitness for cell division. Cells lacking IRS2 were found to have a shortened mitotic phase only when exposed to mitotic stress, suggesting a role for IRS2 in the regulation of the mitotic checkpoint, the spindle assembly checkpoint (SAC), which monitors the microtubule engagements by kinetochores to ensure the accurate segregation of chromosomes to maintain genome integrity ^10, 11^. Although this study implicates IRS2 in protection from mitotic stress, the mechanism by which IRS2 contributes to this mitotic regulation has not been established.

In the current study, we investigated IRS2 expression and regulation during mitosis to understand how IRS2 functions in the cellular response to mitotic stress. Our studies reveal a role for IRS2 in PLK1-dependent mitotic regulation that is independent of its function as a signaling adaptor for the IIS pathway.

## Results

### IRS2 expression and phosphorylation are regulated during cell cycle progression

To study the mechanistic role of IRS2 in mitosis, IRS2 expression during cell cycle progression was assessed in the human triple negative breast cancer (TNBC) cell lines SUM-159 and MDA-MB-231. TNBC models were used because they express elevated levels of IRS2 and they are characterized by a high proliferation rate and chromosome instability, which requires an active mitotic checkpoint to maintain viability ^12–14^. Cells were synchronized using a sequential thymidine-nocodazole block and analyzed for cell cycle stage by flow cytometry and IRS2 abundance by immunoblot (Fig 1A). IRS2 expression was low in thymidine arrested cells (G_1_/S), increased when released from thymidine for 8 hrs (G_2_/M), remained elevated after nocodazole treatment to arrest in mitosis (M) and decreased after nocodazole release into normal growth media (return to G_1_). Similar expression changes were observed after palbociclib arrest (G_1_) and release (Fig S1A). Increased IRS2 protein expression in G_2_/M was also observed using flow cytometry to co-stain for IRS2 and DNA (Fig S1B). Analysis of mRNA expression under the same treatment conditions revealed that the observed changes in IRS2 protein expression in G_2_/M was not dependent upon transcriptional increases, because IRS2 mRNA expression modestly decreased in G_2_/M arrested cells (Fig S1C). This result likely reflects a global reduction in mRNA transcription during mitosis that occurs due to chromatin condensation and transcription silencing^15^. Upon exit from mitosis, IRS2 mRNA expression increased, which may reflect the requirement to restore IRS2 protein levels for signaling functions during interphase.

**Figure 1.**
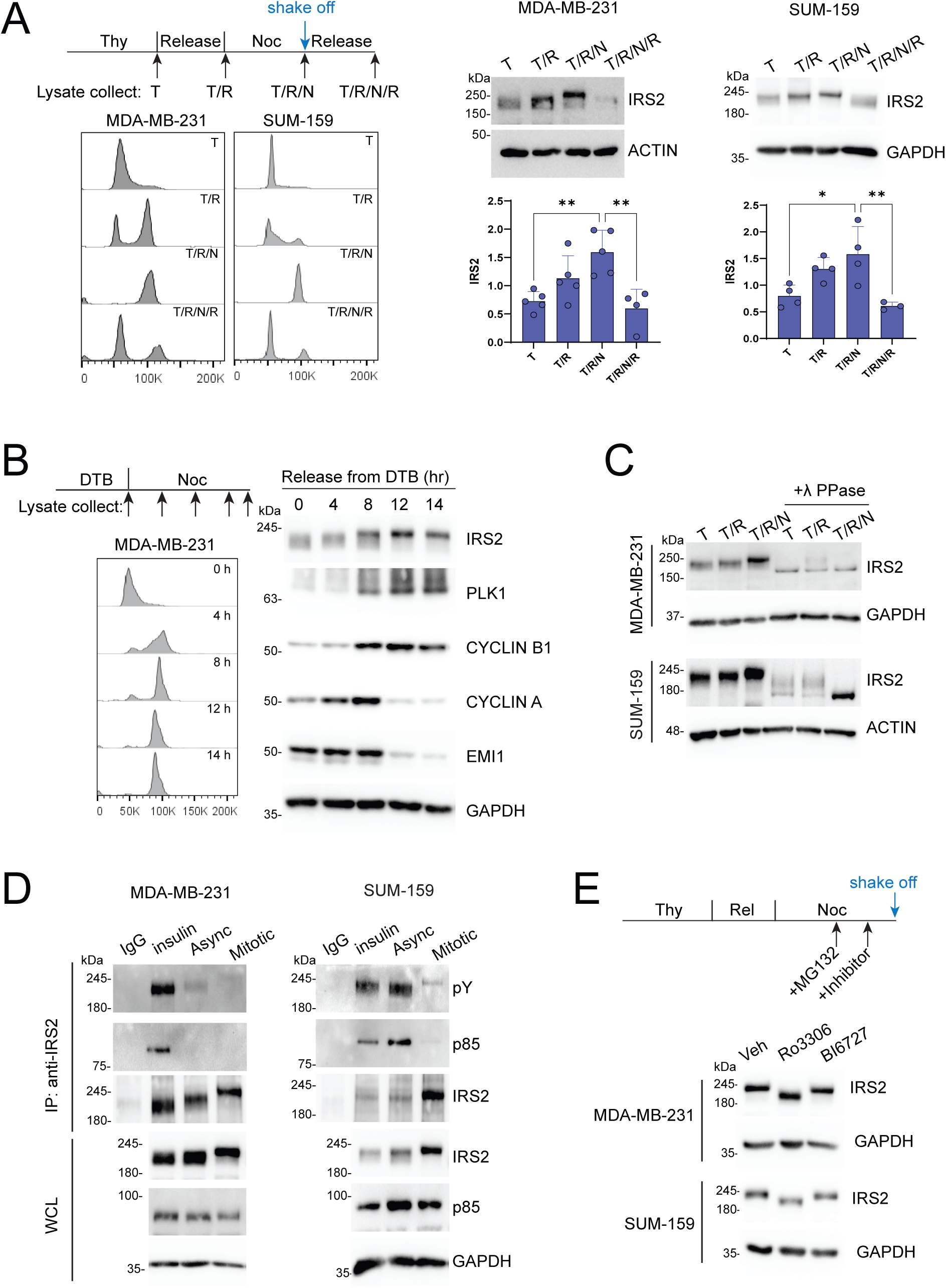
(A) MDA-MB-231 and SUM-159 cells were treated with thymidine (2 mM) for 18 hrs (T), released for 8 hrs into normal growth media (T/R), then treated with nocodazole (90 ng/ml) for 16 hrs. Mitotic cells were collected by shake-off (T/R/N) and then replated and released into growth media for 5 hrs (T/R/N/R). Cell cycle stages were analyzed by flow cytometry (left) and IRS2 expression was analyzed by immunoblotting (right). The data shown in the graphs represent the mean ± SD of three or more independent experiments. (B) MDA-MB-231 cells were synchronized by double thymidine block (DTB) and released into growth media in the presence of nocodazole (90 ng/ml). Cell cycle stages were analyzed by flow cytometry (left) and cell extracts were analyzed by immunoblotting for IRS2 and cell cycle proteins. (C) MDA-MB-231 and SUM-159 cells were treated as shown in (A) and cell extracts were left untreated or treated with λ phosphatase before analysis by immunoblotting. (D) Cells were serum starved overnight and stimulated with insulin (100 ng/ml) for 10 min (insulin), grown asynchronously in normal growth media (Async), or arrested in mitosis by treatment with nocodazole (90 ng/ml, 16 hrs) and shake-off (Mitotic). Cell extracts were immunoprecipitated and analyzed by immunoblotting. (E) MDA-MB-231 and SUM-159 cells were treated with thymidine for 16 hrs, released for 8 hrs, and then mitotic cells were enriched by nocodazole treatment (90 ng/ml, 16 hrs). 1 hour before mitotic cell collection, cells were treated with MG132 (10 uM) and then treated with Ro3306 (10 uM) or BI6727 (100 nM) for the final 30 min before cell collection. Cell extracts were isolated and analyzed by immunoblotting as labeled. All immunoblots and flow plots are from representative experiments that were replicated at least three times independently.

Cell cycle progression is a tightly orchestrated process that is controlled by protein degradation mediated by two families of E3 ubiquitin ligases, the APC/C and the Skp1-Cul1-F-box protein (SCF) complex ^16^. APC/C activity is temporally regulated through the combination of two coactivators, CDC20 (required for mitotic progression) and CDH1 (required for G_1_ stabilization). IRS2 was identified as an APC/C^CDH1^-specific substrate that is degraded as cells exit mitosis and re-enter G_1_ ^8^. To evaluate more rigorously how IRS2 expression fluctuates during cell cycle progression, cells were arrested at the G_1_/S boundary with a double thymidine block and then released into normal growth medium containing nocodazole to allow cells to progress through S and G_2_ and arrest in M. Cell cycle phases were monitored by flow cytometry and the expression of known cell cycle proteins (Figs 1B and S1D). Early mitotic inhibitor (EMI1) inhibits both APC/C^CDC20^ and APC/C^CDH1^ during S phase and EMI1 expression is targeted for degradation in early mitosis by the SCF complex ^17^. EMI1 expression was high at G_1_/S and decreased after 8 hrs as cells progressed into G_2_/M. Expression of CYCLIN A gradually increased until 8 hrs after release and then also decreased at later time points. This pattern reflects prior observations indicating stable CYCLIN A during S/G_2_ and the initiation of degradation immediately after nuclear envelope breakdown facilitated by APC/C^CDC20^ regardless of SAC activation ^18–20^. CYCLIN B1 expression increased within 8 hrs as cells entered into G_2_/M and remained high in cells arrested in mitosis in the presence of nocodazole, reflecting that SAC activation delays its degradation ^19^. Expression of the mitotic kinase Polo like kinase 1 (PLK1) also increased within 8 hrs and remained elevated in nocodazole because of the delayed mitotic exit ^21^. The expression pattern of IRS2 was similar to CYCLIN B1 and PLK1, peaking in M and remaining high during delayed mitotic exit.

In addition to cell cycle phase-dependent changes in IRS2 protein expression levels, IRS2 mobility by SDS-PAGE is reduced in mitotic cells (Fig 1A and 1B). This IRS2 mobility shift has been previously reported and suggested to be attributed to phosphorylation during mitosis ^8^. To directly assess the impact of phosphorylation on the mobility shift of IRS2, protein lysates were treated with λ phosphatase *in vitro* prior to analysis by SDS-PAGE and immunoblot. After treatment with λ phosphatase, IRS2 mobility was increased across all cell cycle phases, indicating that phosphorylation is responsible for the shift in mobility observed in mitosis (Fig 1C). In its role as an adaptor protein for the IIS pathway, IRS2 is phosphorylated on multiple tyrosine residues by the intrinsic insulin and IGF-1 receptor kinases ^2, 22^. To determine if the mobility shift of IRS2 that is observed during mitosis is related to tyrosine phosphorylation of the protein, IRS2 was immunoprecipitated from extracts of insulin stimulated cells, asynchronously growing cells, or mitotic cells. The immune complexes were immunoblotted with antibodies specific for phosphotyrosine or the p85 regulatory subunit of PI3K, which is recruited to IRS2 in a phosphotyrosine-dependent manner. Little to no IRS2 tyrosine phosphorylation or recruitment of p85 was observed in mitotic cells when compared with the levels observed after insulin stimulation (Fig 1D). IRS2 tyrosine phosphorylation and p85 binding were high in asynchronously growing SUM-159 cells relative to MDA-MB-231 cells, which is attributed to the inclusion of insulin in the normal growth medium of SUM-159 cells. Therefore, increased tyrosine phosphorylation is not the underlying cause of the IRS2 mobility shift in mitosis.

We next assessed the contribution of mitotic regulatory kinases to the mobility shift of IRS2 during mitosis. The serine/threonine kinase PLK1 has been previously reported to directly phosphorylate IRS2 ^23^. In addition, inhibition of CDK1, which regulates the transition from G_2_ to M, prevents the mobility shift of IRS2 ^8^. However, the inhibition of CDK1 also arrests cells in G_2_ and prevents entrance into mitosis, which could indirectly inhibit the phosphorylation of IRS2. To directly investigate the involvement of PLK1 and CDK1 in the mitosis-dependent phosphorylation of IRS2, cells were synchronized and arrested in mitosis with nocodazole treatment for 16 hrs before treatment with MG132 to prevent proteasomal degradation. CDK1 and PLK1 inhibitors were added for 1 hr before collecting mitotic cells and analysis by immunoblotting (Fig 1E). The CDK1-selective inhibitor Ro3306 reduced the mobility shift of IRS2, whereas the PLK1-specific inhibitor BI6727 did not alter IRS2 mobility significantly, when compared with vehicle control. We conclude from these data that the majority of IRS2 phosphorylation that occurs during mitosis can be primarily attributed to the activity of CDK1.

### IRS2 sustains SAC-dependent mitotic arrest in response to mitotic stress

Depletion of IRS2 shortens mitotic duration in cells exposed to mitotic stress, a finding that implicates IRS2 in the regulation of the SAC ^8, 24^. This checkpoint is essential for correct chromosome segregation in mitosis by integrating signals from microtubule-unattached kinetochores and delaying anaphase onset until chromosomes are fully attached ^10^. We used SUM-159 and MDA-MB-231 cells that were genetically modified by CRISPR/Cas9 gene editing to disrupt IRS2 expression (IRS2^-/-^) to investigate the contribution of IRS2 to the regulation of mitotic arrest in response to mitotic stress (Fig S2A). In asynchronous culture conditions, in the absence of mitotic stress, cell cycle profiles were similar in non-targeting (sgNT) control and IRS2^-/-^ cells (Fig. S2B). Cells were transfected to express mCherry-Histone-H2B to monitor DNA content and exposed to nocodazole for 10 hrs to induce mitotic stress and activate the SAC ^25^. The number of mitotic cells was assessed by imaging of nuclear morphology (Fig S2C). Fewer mitotic cells were present in the IRS2^-/-^ population relative to cells expressing IRS2 (sgNT) after 10 hrs of nocodazole treatment (Fig 2A)^8^.

**Figure 2.**
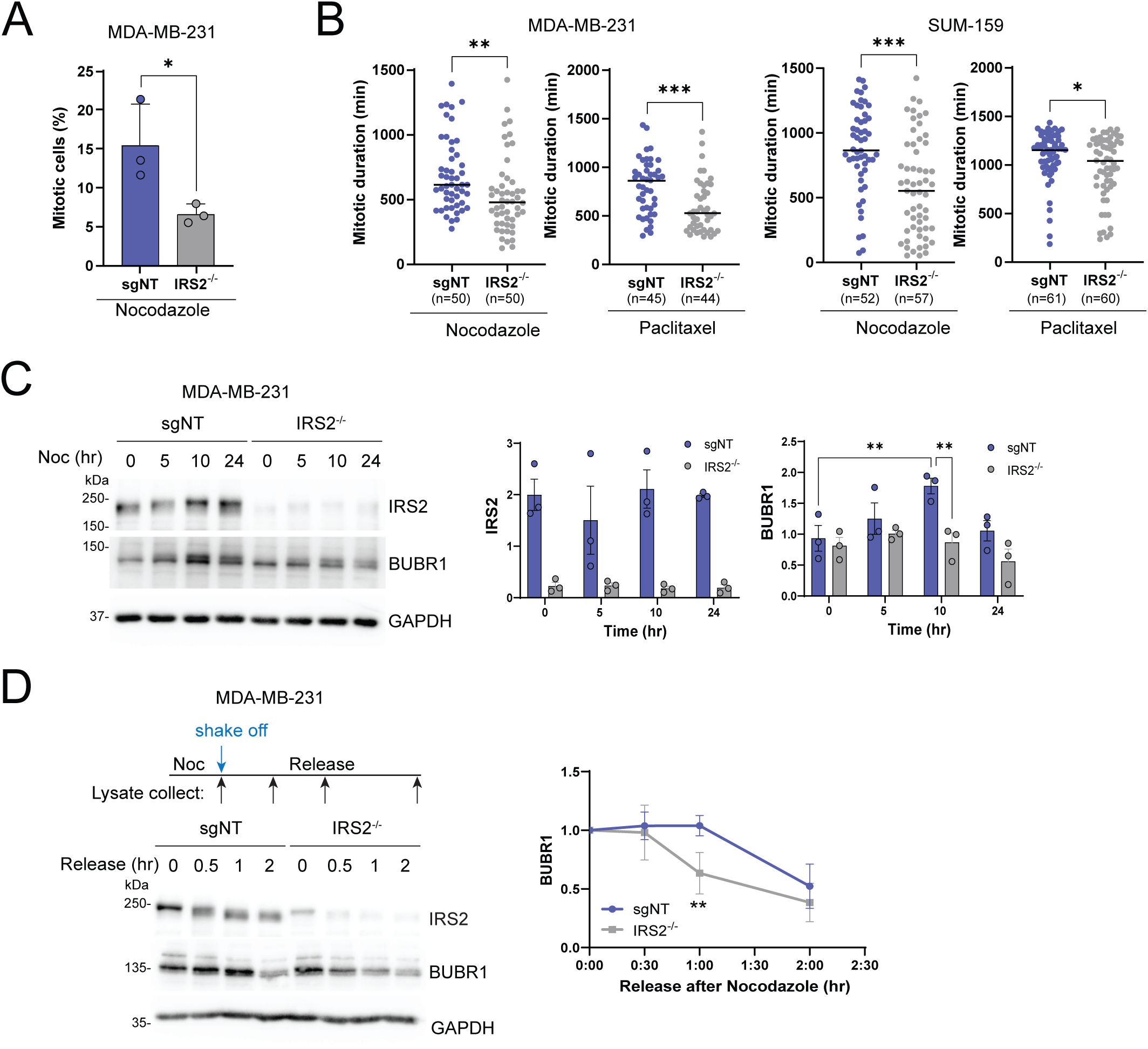
(A) H2B-mCherry labeled MDA-MB-231 cells treated with non-targeting guide RNA (*sgNT*) or IRS2 knockout (KO) cells (*IRS2^-/-^*) were treated with nocodazole (90 ng/ml). Cells were imaged at 10 hours after the treatment, followed by analysis of mitotic cells. The data shown represent the mean ± SD of a representative experiment performed three times independently. (B) H2B-mCherry labeled MDA-MB-231 and SUM-159 cells were treated with nocodazole (90 ng/ml) or paclitaxel (100 nM) and imaged every 10 minutes by widefield time-lapse microscopy for 24 hrs. Each dot represents the mitotic duration of an individual cell measured as the time from nuclear envelope breakdown to division, slippage, cell death or imaging termination (arrest). The data shown represent the mean ± SD of a representative experiment performed three times independently. (C) sgNT or *IRS2^-/-^* MDA-MB-231 cells were treated with nocodazole (90 ng/ml) for the indicated times. Lysates were isolated from both adherent and nonadherent cells and analyzed by immunoblotting as labeled. The data shown in the graphs represent the mean ± SD of three independent experiments. (D) *sgNT* or *IRS2^-/-^* MDA-MB-231 cells were treated with nocodazole (90 ng/ml) for 16 hrs, mitotic cells were collected by shake-off then released into growth media for the indicated times. Cell extracts were isolated and analyzed by immunoblotting as labeled. The data shown in the graph on the right represent the mean ± SD of three independent experiments.

To investigate further the impact of IRS2 on mitotic duration in response to mitotic stress, time-lapse imaging was used to follow mitotic progression in the presence of nocodazole or the microtubule stabilizing drug paclitaxel. The time from nuclear envelope breakdown (NEB) to anaphase onset was measured in each cell population to monitor mitotic progression. As reported previously, loss of IRS2 did not alter mitotic duration in normal growth conditions (Fig. S2D). In contrast, in both MDA-MB-231 and SUM-159 cells, IRS2^-/-^ cells displayed shortened mitotic duration under both nocodazole and paclitaxel treatment conditions (Figs 2B and S2E). Importantly, the time to mitosis entry did not show a significant difference under any of the treatment conditions (Fig S2F), providing further evidence that the decreased mitotic duration observed in the absence of IRS2 expression in response to mitotic stress is attributed to early mitotic exit and not delayed mitotic entry. SAC arrest is mediated through the mitotic checkpoint complex (MCC), which consists of four proteins; MAD2, BUBR1, BUB3 and CDC20. BUBR1 is phosphorylated and acetylated during mitosis for stabilization, and its degradation signals SAC silencing ^26, 27^. In MDA-MB-231 cells treated with nocodazole, BUBR1 expression increased initially and then decreased with prolonged drug treatment, reflecting SAC activation (10 hrs) and partial mitotic exit (24 hrs) (Fig 2C). In contrast, BUBR1 expression did not accumulate at the 10 hr timepoint in IRS2^-/-^ cells, indicating a weak SAC. Cells arrested in mitosis by nocodazole treatment were collected and replated into normal growth medium to evaluate their rate of mitotic exit. BUBR1 expression decreased more rapidly in IRS2^-/-^ cells than in sgNT (Fig 2D). Taken together, our data support that cells lacking IRS2 elicit a weakened SAC in response to mitotic stress that results in faster mitotic exit.

### IRS2-regulated mitotic arrest prevents mitotic catastrophe

Activation of the SAC arrests cells in prometaphase until all kinetochores have attached properly to the mitotic spindle and chromosomes can be separated with fidelity. If mitotic stress is prolonged and the SAC is functionally deficient, cells may exit mitosis before complete alignment of the chromosomes, leading to chromosomal defects and either aneuploidy or cell death. Indeed, 30hrs incubation in nocodazole led to an increase in cells exhibiting nuclear fragmentation, an indication of cell death (Fig 3A and S2C). To determine if loss of IRS2 expression resulted in abnormal chromosome segregation at mitotic exit, two independent sgNT and IRS2^-/-^ SUM-159 subclones were treated with nocodazole (16 hr) to arrest cells at prometaphase and then released into normal culture media to examine cells as they progress through anaphase. In the absence of IRS2, significantly more cells were observed undergoing abnormal chromosome segregation which included lagging chromosomes, chromosome bridges and multipolar segregation (Fig 3B). To determine the consequence of IRS2 loss on cellular outcomes in response to prolonged mitotic stress, cell fates from the mitotic duration experiment in Fig 2B were characterized into four categories (Fig 3C): 1) Arrest: cells remained in mitosis at the endpoint of imaging; 2) Mitotic death: cells underwent apoptosis right after mitotic exit; 3) Slippage: cells exited mitosis without proper division; and 4) Division: cells exited mitosis, producing two independent daughter cells. Cell fate outcomes differed between the SUM-159 and MDA-MB-231 sgNT control cells, with the majority of SUM-159 cells arresting in response to drug treatment and the majority of MDA-MB-231 cells exhibiting slippage. However, both cell lines underwent increased cell death when IRS2 expression was lacking (Fig 3D). Collectively these observations support that IRS2 protects cells from cell death in response to mitotic stress by supporting a stable SAC and preventing mitotic catastrophe.

**Figure 3.**
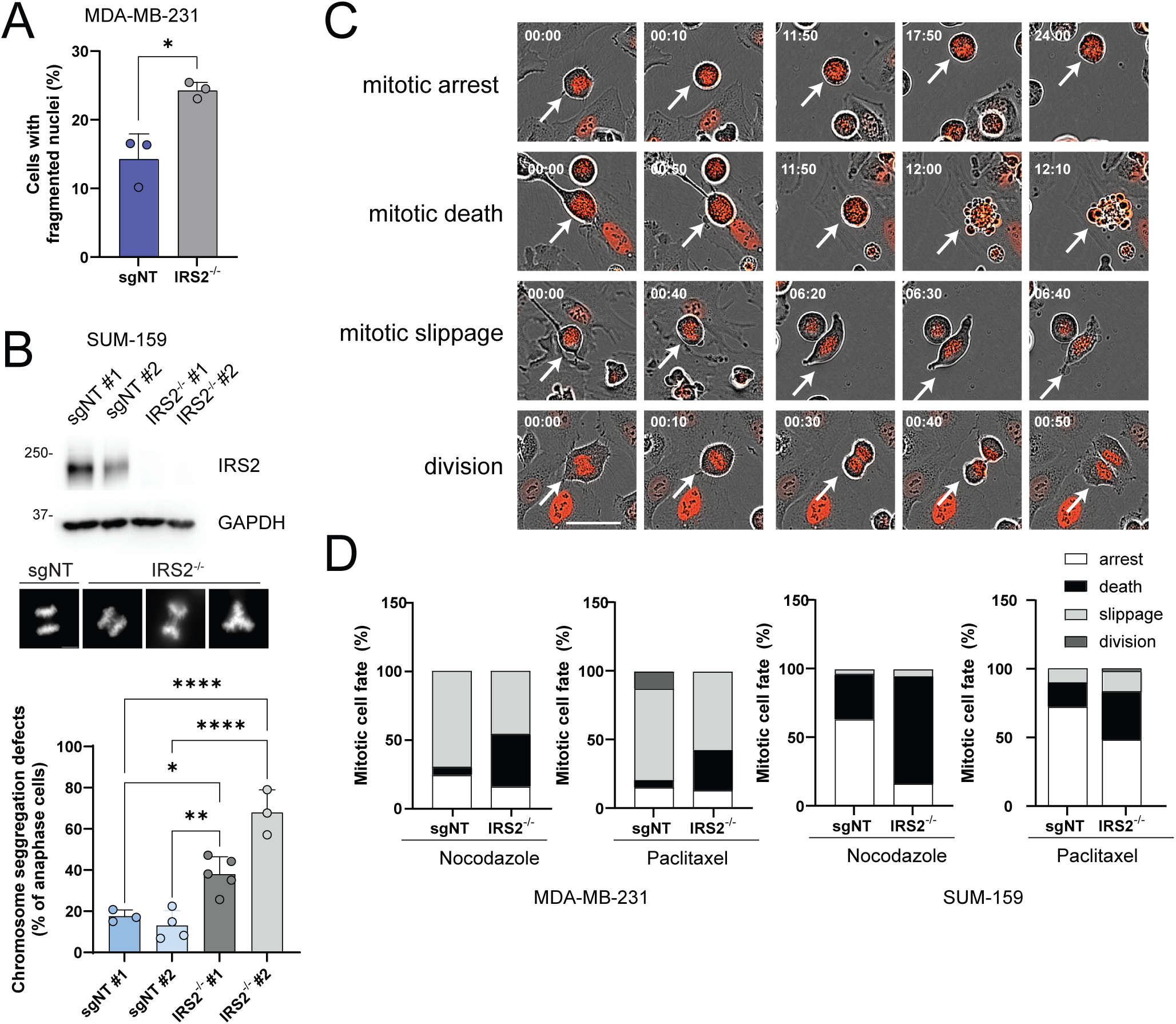
(A) H2B-mCherry labeled MDA-MB-231 cells treated with non-targeting guide RNA (*sgNT*) or IRS2 knockout (KO) cells (*IRS2^-/-^*) were treated with nocodazole (90 ng/ml). Cells were imaged at 30 hours after the treatment, followed by analysis of fragmented nuclei. (B) SUM-159 single cell sgNT and IRS2^-/-^ clones were generated and expression of IRS2 was assessed by immunoblot. Cells were treated with nocodazole (50 ng/ml) for 16 hrs and then released into growth media for 45 min. Cells were fixed and stained with Hoechst33342 and anaphase cells were imaged. Chromosomal segregation defects were quantified (representative images shown for lagging chromosomes, chromosome bridges and multipolar segregation). The data shown represent the mean ± SD of a representative experiment performed three times independently. (C and D) Representative images (C) and quantification of cell fates (D) for each cell line and drug. Scale bar, 50 μm. Time=hr:min. For (D), MDA-MB 231 sgNT, nocodazole, n=50; MDA-MB-231 IRS2^-/-^, nocodazole, n=50; MDA-MB 231 sgNT, paclitaxel, n=45; MDA-MB-231 IRS2^-/-^, paclitaxel, n=44; SUM-159 sgNT, nocodazole, n=52; SUM159 IRS2^-/-^, nocodazole, n=57; SUM-159 sgNT, paclitaxel, n=61; SUM159 IRS2^-/-^, paclitaxel, n=60.

### IRS2 interacts with PLK1 to regulate mitotic progression

As an adapter protein, IRS2 mediates its functions through binding partners. To further investigate the mechanism of action of IRS2 in mitosis, we analyzed data from an IRS2 interactome study we had previously performed. In this study, IRS1^-/-^/IRS2^-/-^ SUM-159 cells expressing FLAG-tagged IRS2 were serum starved and then stimulated for 15 min with insulin, after which IRS2 was immunopurified from cell extracts with FLAG-specific antibody-conjugated agarose beads. The immune complexes were eluted with FLAG peptides, the eluates were trypsin-digested and peptides were analyzed by mass-spectrometry. This analysis identified 42 proteins with statistically increased binding to IRS2 in response to insulin stimulation, including the p85 regulatory subunit of PI3K, a known IRS2 interacting protein that served as an internal control for signaling activation (Fig 4A and Table S1). Although this interactome study had not been designed to enrich for mitotic cells, we hypothesized that a small number of cells in this asynchronous population were likely in the mitotic state. Analysis of our dataset revealed a significant interaction of IRS2 with only one known mitotic regulator, PLK1, a finding that had been previously reported ^23, 28, 29^. We also interrogated an open access database, the Biological General Repository for Interaction Datasets (Biogrid), ^30^ to determine if additional cell cycle regulatory proteins that could mediate the mitotic functions of IRS2 had been reported. No additional proteins with known roles in mitotic regulation were identified through this analysis.

**Figure 4.**
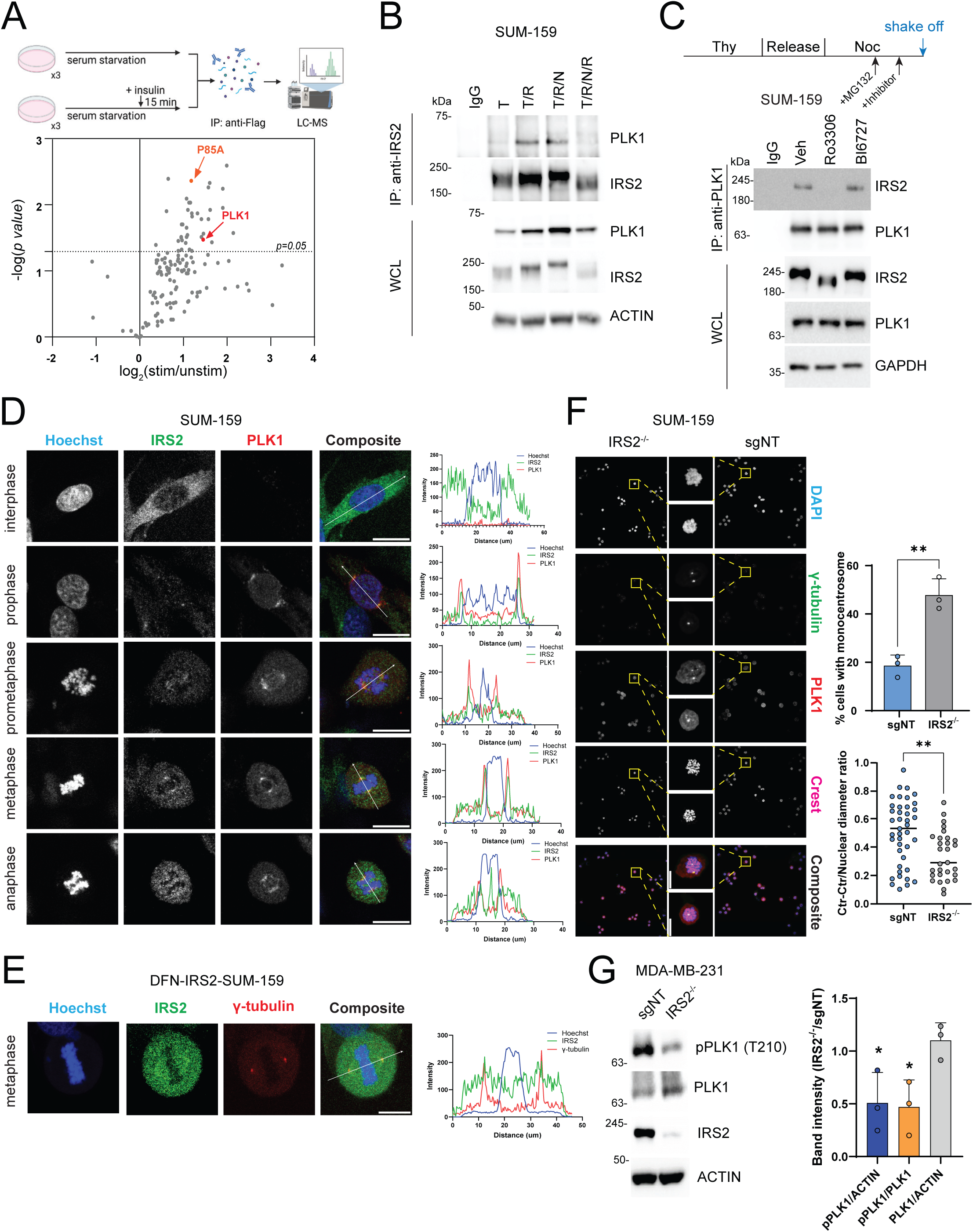
(A) *Top,* schematic of experimental process for mass spectrometry analysis. IRS1^-/-^/IRS2^-/-^ SUM159 cells expressing FLAG-tagged IRS2-WT were serum starved and then stimulated with insulin (500 ng/ml) for 15 min. Cell extracts were immunoprecipitated with anti-FLAG antibodies followed by LC/MS analysis. *Bottom,* Volcano plot comparing proteomes of unstimulated vs stimulated IRS2-WT expressing cells. (B) SUM-159 cells were synchronized as shown in Fig 1A and cell extracts were immunoprecipitated and immunoblotted as labeled. (C) SUM-159 cells were synchronized as shown in the schematic and cell extracts were immunoprecipitated and immunoblotted as labeled. (D) SUM-159 cells were grown asynchronously (interphase) or arrested at the G_2_/M boundary by Ro3306 (10 uM) treatment for 20 hrs and then released into growth media for 30 min. Plots on the right represent the line scan analysis of respective composite images. White arrows depict the area used for the line scan analysis. Scale bar, 20 μm. (E) DFN-IRS2 SUM-159 cells were treated as in Fig 4D and imaged for IRS2 and γ-tubulin. Scale bar, 20 μm. (F) SUM-159 cells (sgNT and IRS2^-/-^) were arrested in prometaphase by nocodazole treatment (50 ng/ml, 16 hrs), followed by immunostaining, scale bar, 10 um. (Top graph) Quantification of cells with monocentrosomes determined by γ-tubulin staining. sgNT, n=50; IRS2^-/-^, n=53. (Bottom graph) Quantification of distance between two centrosomes (normalized to nuclear diameter). sgNT, n=41; IRS2^-/-^, n=29. (G) MDA-MB-231 cells (sgNT and IRS2^-/-^) were treated with nocodazole (90 ng/ml) for 16 hrs and cell extracts from mitotic cells were isolated and analyzed by immunoblotting (left). Densitometry analysis from three independent experiments (right). All immunoblots are from representative experiments that were replicated at least three times independently.

PLK1 plays a multifaceted role in mitosis, including the regulation of mitotic entry, centrosome maturation and separation, spindle formation, SAC activation and cytokinesis ^31^. To explore the functional significance of the IRS2-PLK1 interaction, we confirmed that endogenous IRS2 and PLK1 interact in mitotic SUM-159 and MDA-MB-231 cells (Fig S3A). This interaction is enriched in cells synchronized in the G_2_/M phase of the cell cycle (Fig 4B) and the binding is dependent upon CDK1 activity, but not PLK1 activity (Fig 4C). These findings are consistent with a previous PLK1 proteomic interactome study that reported PLK1 binding to IRS2 requires CDK1 activity, but not PLK1 activity ^28^. We next examined the intracellular localization of IRS2 during mitosis to assess its co-localization with PLK1. PLK1 predominantly localizes at centrosomes and kinetochores during prophase, prometaphase and metaphase and translocates to the spindle midzone as chromosome segregation occurs in anaphase ^31, 32^. IRS2 localization during mitosis is less well understood, with only one report that IRS2 phosphorylated on pS1098 (mouse sequence) localizes to the spindle poles during metaphase ^23^. SUM-159 cells were cultured in either normal growth media (predominantly interphase cells) or synchronized at the G_2_/M boundary by treatment with the CDK1 inhibitor (Ro3306) for 20 hrs and then released into growth media for 30 min to enrich for mitotic cells before fixation and immunostaining for IRS2 and PLK1 (Figs 4D and S3B). During interphase, IRS2 is predominantly localized in the cytosol and PLK1 is not expressed (Fig 4D) ^24^. In prophase, IRS2 is enriched and co-localizes with PLK1 at centrosomes and this co-localization persists during prometaphase (see intensity plots on right). In metaphase, IRS2 and PLK1 are both enriched at the spindle poles, exhibiting their strongest co-localization. As cells progress to anaphase, PLK1 remains at the spindle plate, but a specific co-enrichment of IRS2 is no longer observed. To further validate the localization of IRS2 during mitosis, we imaged SUM-159 cells in which an mNeonGreen tag was inserted into the N-terminus of the endogenous IRS2 locus by CRISPR-Cas9 gene targeting (DFN-IRS2) ^33^. As we previously reported, IRS2 is expressed diffusely in the cytoplasm during interphase in these cells (Fig S3C). After synchronization of cells at G_2_/M with Ro3306 and then release into normal growth medium, endogenous IRS2 co-localized with γ-tubulin, a marker of centrosomes, during metaphase (Fig 4E). An enhanced IRS2 localization along the mitotic spindle was also observed in these endogenously tagged cells.

To determine if PLK1 binding to IRS2 determines its localization at the spindle pole, sgNT and IRS2^-/-^ SUM-159 cells were arrested in prometaphase with nocodazole and stained for PLK1 and γ-tubulin (Fig 4F). PLK1 co-localized with γ-tubulin independently of IRS2 expression, indicating that the interaction between PLK1 and IRS2 is not required for the centrosomal localization of PLK1. However, a 2-fold increase in cells exhibiting monocentrosomes was observed in cells lacking IRS2 expression. Moreover, when two centrosomes were present in IRS2^-/-^ cells, the distance between the centrosomes was significantly reduced. The fluorescence intensity of γ-tubulin staining at monocentrosomes was higher than the staining intensity at double centrosomes, supporting that IRS2 is not required for centrosome duplication (Fig S3D). Impaired centrosome separation can lead to defects in bipolar spindle formation and abnormal chromosome segregation, which is consistent with the increased chromosome segregation defects we observed in cells depleted of IRS2 (Fig 3B).

Cells with reduced PLK1 expression or activity are also characterized by monocentrosomes and increased chromosomal abnormalities ^34^. Despite PLK1 localizing normally to centrosomes in IRS2^-/-^ cells, the reduction in centrosome separation in these cells raised the possibility that PLK1 activity could be impaired in the absence of IRS2. Phosphorylation of T210 in the kinase domain of PLK1 is essential for PLK1 activation and can be used to monitor PLK1 activity ^35, 36^. The phosphorylation of T210 is initiated by Aurora A-Bora complex in G_2_ and serves a role in G_2_/M transition ^37, 38^, but how this phosphorylation is maintained and enhanced during mitosis is still not clear. To determine if IRS2 regulates PLK1 activity, mitotic cells were collected and immunoblotted for pT210-PLK1. A significant reduction in PLK1 phosphorylation was observed in IRS2^-/-^ cells relative to sgNT control cells (Fig 4G). Our data suggests that while PLK1 is phosphorylated on T210 to enable G_2_/M transition in IRS2^-/-^ cells, IRS2 sustains or enhances this phosphorylation during mitosis. In its absence, PLK1 does not achieve or maintain full activation and centrosome separation is impaired.

We sought to identify the residues of IRS2 that are responsible for the recruitment of PLK1 to investigate the functional significance of the interaction of IRS2 with PLK1 for mitotic regulation. PLK1 consists of two functional domains: the kinase domain (KD) in the N-terminal region and the polo-box domain (PBD) in the C-terminal region. The PBD specifically recognizes PBD motifs (Ser-pSer/pThr-Pro) present in its binding partners ^39^. Analysis of the IRS2 sequence revealed eight potential PBD motifs (Fig 5A), which are conserved across mammals (Fig 5B). Initially we compared PLK1 binding of IRS2-WT with an IRS2 mutant (IRS2Δ846) that lacks the three C-terminal PBD motifs (S953, S985, S1012). Cells were synchronized with thymidine and nocodazole treatment and the ability of IRS2 to co-immunoprecipitate PLK1 was evaluated. IRS2-WT bound PLK1 in cells in the G_2_/M phase (T/R) and this binding increased in cells enriched in mitosis (T/R/N). The IRS2Δ846 mutant retained the ability to bind to PLK1 at an equivalent level when compared with IRS2-WT (Fig S4A), indicating that the C-terminal PBD domains are not required for PLK1 binding.

**Figure 5.**
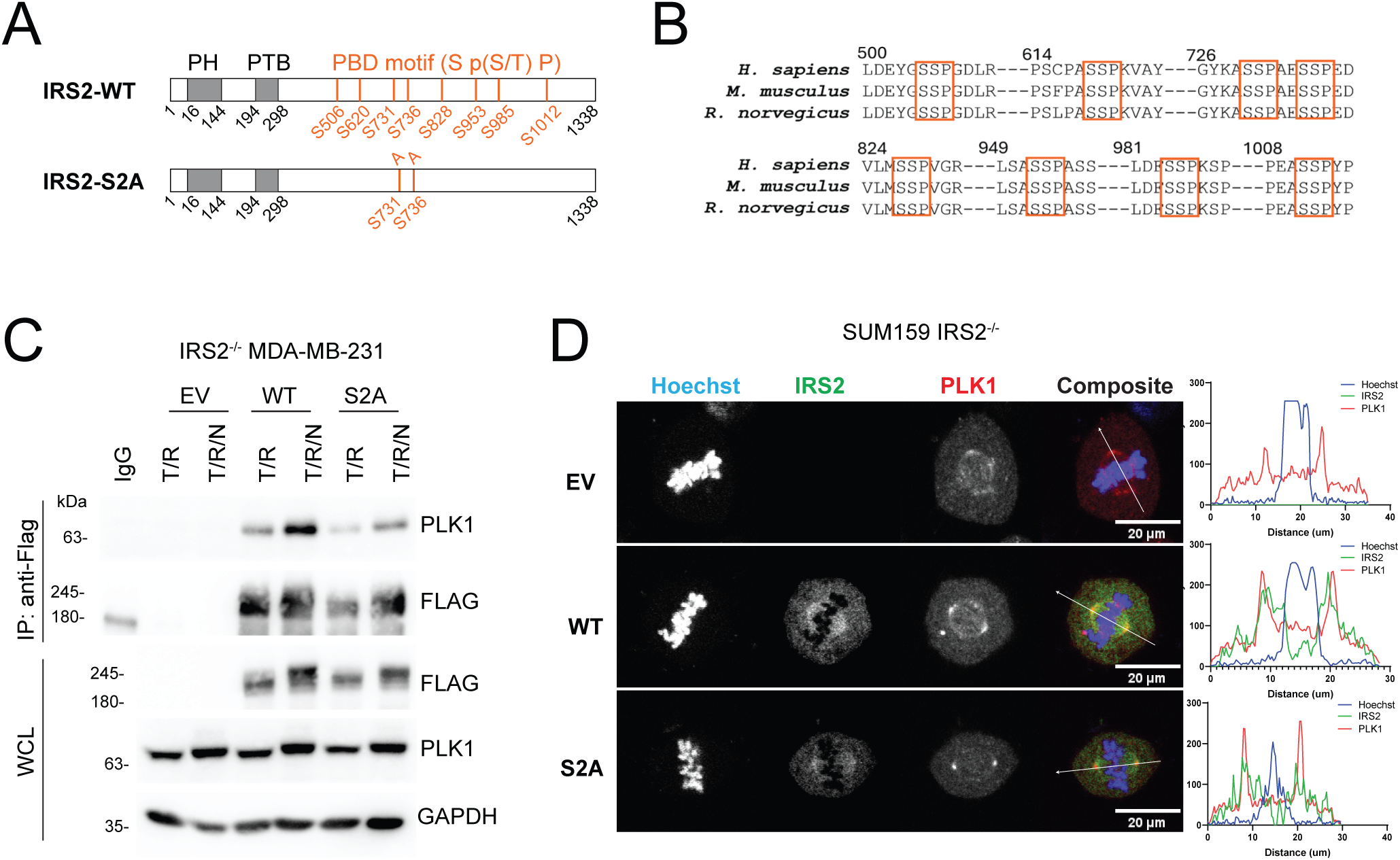
(A) Schematic showing IRS2-WT and IRS2-S2A phospho-mutant. PH, pleckstrin homology domain; PTB, phosphotyrosine binding domain; PBD, polo-box domain. (B) IRS2 sequence alignment showing that the PBD motifs (red boxes) are conserved across mammals. (C) *IRS2^-/-^* MDA-MB-231 cells expressing EV, IRS2WT or IRS2-S2A were synchronized by thymidine/release (T/R; G_2_/M) or thymidine/release/nocodazole and shake-off (T/R/N; M). Cell extracts were immunoprecipitated with FLAG-specific antibodies and immunoblotted as labeled. (D) *IRS2^-/-^* SUM-159 cells expressing EV, IRS2-WT or IRS2-S2A were arrested at the G_2_/M boundary by Ro3306 (10 uM) treatment for 20 hrs and then released into mitosis for 30 min. Cells were fixed and processed for immunofluorescence. Scale bar, 20 μm.

We next examined our LC/MS dataset and determined that of the remaining canonical PBD motifs, only S731 and S736 were phosphorylated. We generated an IRS2 mutant in which both S731 and S736 were mutated to alanine (S731A and S736A; IRS2-S2A) and expressed this mutant in IRS2^-/-^ MDA-MB-231 cells. Binding of PLK1 to the IRS2-S2A mutant was markedly diminished in both G_2_/M and M when compared with IRS2-WT, confirming that S731 and S736 are involved in PLK1 recruitment to IRS2 (Fig 5C). Given that our LC/MS analysis was performed on cells primarily in interphase, the low residual binding of IRS2-S2A to PLK1 may be due to additional sites that are phosphorylated in a mitosis-specific manner that were not identified in our analysis. The observation that IRS2-S2A still exhibits a mobility shift in mitosis also reflects that additional phosphorylation sites are present. Although we did not observe significant tyrosine phosphorylation of IRS2 in mitotic cells (Fig 1E), we evaluated the binding of PLK1 to an IRS2 mutant in which five tyrosine residues (Y542, Y653, Y675, Y742 and Y823) located within canonical PI3K binding motifs were substituted with phenylalanine (IRS2-Y5F). Tyrosine phosphorylation of this IRS2 mutant is significantly diminished and it is deficient in its recruitment of PI3K (p85) (Fig S4B). As a result, cells expressing this mutant exhibit reduced glucose uptake, invade less well and have diminished stemness in response to insulin/IGF signaling ^5, 7, 22^. The IRS2-Y5F mutant maintained the ability to interact with PLK1 at WT levels during mitosis (Fig S4C).

Cells expressing IRS2-WT or IRS2-S2A were imaged in metaphase to evaluate their co-localization with PLK1. PLK1 expression was enriched at the spindle poles in cells expressing both IRS2-WT and IRS2-S2A. Both IRS2-WT and IRS2-S2A co-localized with PLK1 at the spindle poles, (Fig 5D), indicating that targeting IRS2 to the spindle poles is not dependent upon the interaction of PLK1 with pS731 and pS736. IRS2-WT and IRS2-S2A were also enriched along the mitotic spindle, as we had observed for endogenously-tagged IRS2 (Fig 4E).

MDA-MB-231 IRS2^-/-^ cells restored with empty vector (EV), IRS2-WT, IRS2-S2A or IRS2-Y5F were evaluated for their response to mitotic stress. Restoration of IRS2-WT and IRS2-Y5F expression increased the number of mitotic cells relative to EV expressing cells after exposure of cells to either nocodazole or paclitaxel for 10 hrs (Fig 6A and S4D). Mitotic duration was also lengthened upon rescue with IRS2-WT (Fig 6B). In contrast, cells expressing the IRS2-S2A mutant exhibited similar mitotic patterns to EV control cells (Figs 6A and 6B). No differences in mitotic entry rate were observed for any of the cell lines (Fig S5A). The contribution of IRS2-PLK1 binding to the regulation of mitotic arrest was further demonstrated by the ability of IRS2-WT, but not IRS2-S2A, to sustain expression of BUBR1 and delay mitotic exit after release from nocodazole arrest (Fig 6C). Finally, cells expressing EV, IRS2-WT or IRS2-S2A were arrested in prometaphase and stained for Hoechst and γ-tubulin. Restoration of IRS2-WT expression to IRS2^-/-^ cells decreased the number of cells exhibiting monocentrosomes and increased the distance of separation between duplicate centrosomes. The ability of IRS2 to interact with PLK1 was necessary for this rescue as centrosome separation in cells expressing the IRS2-S2A mutant was similar to EV expressing cells (Fig 6D).

**Figure 6.**
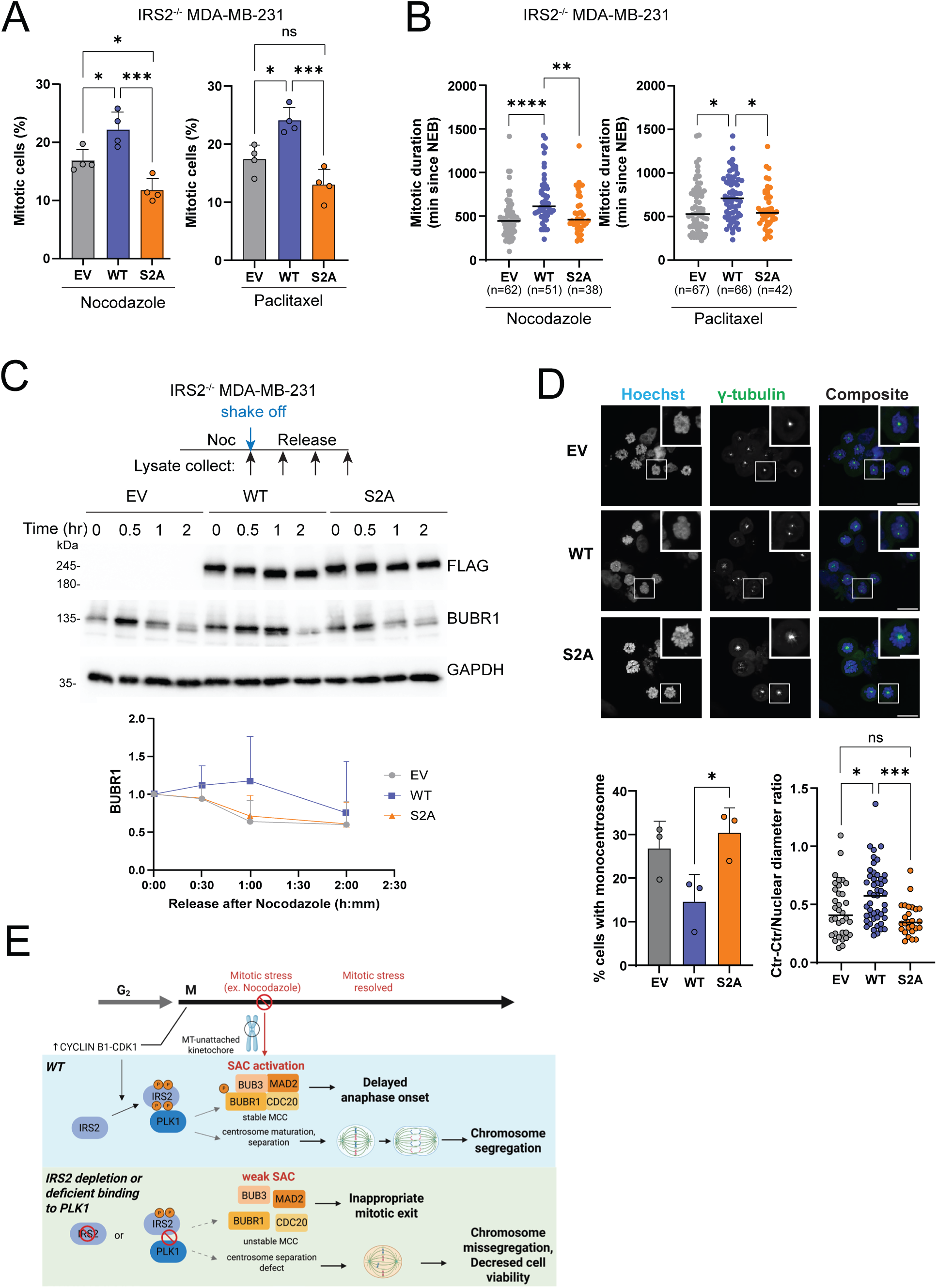
(A) *IRS2^-/-^* MDA-MB-231 cells expressing EV, IRS2-WT or IRS2-S2A were treated with nocodazole (90 ng/ml) or paclitaxel (100 nM). Cells were imaged after 10 hours of treatment. The data shown represent the mean ± SD of a representative experiment performed three times independently. (B) H2B-mCherry labeled *IRS2^-/-^* MDA-MB-231 cells expressing EV, IRS2-WT or IRS2-S2A were treated with nocodazole (90 ng/ml) or paclitaxel (100 nM). Cells were imaged every 10 minutes by widefield time-lapse microscopy for 24 hrs. Each dot represents the mitotic duration of an individual cell measured as the time from nuclear envelope breakdown to division, slippage, cell death or imaging termination (arrest). EV, nocodazole, n=62; WT, nocodazole, n=51; S2A, nocodazole, n=38; EV, paclitaxel, n=67; WT, paclitaxel, n=66; S2A, paclitaxel, n=42. (C) *IRS2^-/-^* MDA-MB-231 cells expressing IRS2-WT or IRS2-S2A were treated with nocodazole (90 ng/ml) for 16 hrs, mitotic cells were collected by shake-off then released into growth media for the indicated times. Cell extracts were isolated and analyzed by immunoblotting as labeled. The data shown in the graph represents the mean ± SD of three independent experiments. (D) *IRS2^-/-^* SUM-159 subclone expressing EV, IRS2WT or IRS2-S2A was arrested in prometaphase by nocodazole treatment (50 ng/ml, 16 hrs), followed by immunostaining (Left) scale bar, 20 um. (Right, top) Quantification of cells with monocentrosomes determined by γ-tubulin staining (left graph). EV, n=73; WT, n=87; S2A, n-=54. Distance between two centrosomes (normalized to nuclear diameter) (right graph). EV, n=34; WT, n=48; S2A, n=26. (E) Schematic of working model.

## Discussion

In this study, we identify a role for the adaptor protein IRS2 in the cellular response to mitotic stress that is distinct from its well-characterized function as a cytoplasmic signaling adaptor. In this latter role, IRS2 is tyrosine phosphorylated by upstream receptor and non-receptor kinases in response to growth factor and cytokine stimuli such as insulin, IGF1 and IL4 to recruit signaling effectors. In contrast, we observe that as cells enter mitosis, IRS2 is phosphorylated on serine/threonine residues, but not tyrosine residues, in a CDK1-dependent manner, and it interacts with and enhances the activity of the cell cycle regulatory kinase PLK1 (Fig 6E). The interaction of IRS2 and PLK1 is important for the regulation of centrosome separation to form a bi-polar spindle and maintenance of the SAC in response to mitotic stress. As a result, in cells that lack IRS2 or express a PLK1-binding deficient mutant, centrosome separation is impaired, and mitotic arrest is weakened when cells are exposed to mitotic stress, leading to chromosomal abnormalities and decreased cell viability. Taken together our results support that IRS2 plays a role in protecting cells from mitotic stress that involves its interaction with PLK1 and these mitotic functions of IRS2 do not involve the canonical signaling functions of this adaptor protein.

An important question that we addressed in our study relates to how the function of IRS2 in mitosis is connected to its role as a signaling adaptor protein during interphase. Previous studies established the importance of IRS2 for the regulation of glucose metabolism, tumor cell migration and invasion and the regulation of MYC and control of breast CSC function by the IIS pathway ^5, 7, 22^. In this capacity, IRS2 is tyrosine phosphorylated by the intrinsic insulin and IGF-1 receptor kinases, which generates binding sites for the regulatory subunit of PI3K, as well as other signaling effectors ^40, 41^. While the majority of studies on IRS2 function have focused on this role as a cytoplasmic signaling adaptor protein during interphase, IRS2 was only recently implicated in regulating the mitotic checkpoint in response to mitotic stress and there was little known about this function ^8^. Our current results reveal that the tyrosine phosphorylation of IRS2 and its recruitment of PI3K are not required for the ability of IRS2 to respond to mitotic stress. This conclusion is based upon the lack of tyrosine phosphorylation of IRS2 during mitosis and the ability of an IRS2 mutant that is deficient in tyrosine phosphorylation and PI3K recruitment retains interactions with PLK1 and to maintain centrosome separation and SAC arrest in response to mitotic stress. These findings indicate that IRS2 has at least two distinct adaptor functions that play important roles in different phases of the cell cycle. However, it should be noted that IRS2 is phosphorylated on serine and threonine residues in a feedback manner after its initial tyrosine phosphorylation, which can modify IRS2 signaling functions and regulate protein stability ^42^. This regulation has been attributed to downstream signaling kinases such as AKT and mTOR/S6K that feedback to control insulin and IGF-1 signaling to maintain growth and metabolic homeostasis. It is possible that this feedback phosphorylation also contributes to the mitotic functions of IRS2 and serves to connect extracellular signals to the control of cell cycle progression.

PLK1 interacts with binding partners through its PBD, which recognizes SpS/TP motifs ^39, 43^ Mutation of two serines in IRS2, S731 and S736, diminished the interaction of IRS2 with PLK1 in G_2_/M. Of note, CDK1 phosphorylates serines or threonines in S/T-P-X-K/R motifs, which does not match the S731 and S736 motif sequences ^39, 43^. The ability of CDK1 inhibitors to prevent PLK1 binding to IRS2 suggests that CDK1 may activate other mitotic kinases to phosphorylate IRS2, or CDK1 phosphorylates IRS2 at other sites to prime the phosphorylation of S731 and S736 by other mitotic kinases. We performed our IRS2 proteomic study using serum starved and insulin stimulated cells (15 min), a cell population that is predominantly in interphase. Given that the interaction of PLK1 with IRS2 was not completely abolished in the IRS2-S2A mutant, there are likely additional mitosis-specific phosphorylation sites in IRS2 that we did not identify. It has previously been reported that PLK1 phosphorylates IRS2 on S560 and S1109 (human sequence) and that these phosphorylation events can impact mitotic duration ^23^. Although we did not detect a significant mobility shift in IRS2 in the presence of PLK1 inhibitors, it is possible that inhibition of only these two phosphosites does not cause an observable shift on an SDS-PAGE gel. However, upon recruitment to IRS2 in response to CDK1 activation, PLK1-dependent phosphorylation of IRS2 could facilitate effector interactions that are important for IRS2 function in mitotic regulation. Additional studies of the IRS2 mitotic interactome will be essential to elucidate the open questions regarding mitotic phosphorylation of IRS2.

We identified two essential mitotic functions that are dependent upon the interaction of IRS2 with PLK1. In early mitosis, IRS2 is required for efficient centrosome separation to form a bipolar spindle. Later in mitosis, the IRS2-PLK1 interaction sustains the SAC to ensure that chromosome segregation is delayed until all kinetochores are engaged by the microtubule spindle. Although spindle assembly can occur without centrosomes, or with impaired centrosome separation due to inhibition of PLK1 activity, bi-polar assembly is inefficient, and this can lead to abnormal chromosome segregation and cell death. In some cell types, such as in the epithelial cells of the larval wing disc of Drosophila melanogaster, the loss of centrosomes alone leads to chromosome segregation abnormalities and increased cell death ^44^. However, if mitosis is delayed, allowing time for bi-polar spindles to form, some cell types can survive in the absence of centrosomes, such as is observed in larval fly brains ^45^. In this instance, a stable SAC is required to delay cell division until an acentrosomal bi-polar spindle can be established ^46, 47^. In the absence of a robust SAC, cells with defective centrosome separation undergo increased cell death, as we have observed in cells in which IRS2 is absent or cannot interact with PLK1. In this role, IRS2 serves to support the fidelity of cell division and sustain cell viability.

Chromosomal instability is a common feature in cancer that provides tumor cells with an adaptive advantage and promotes progression ^48, 49^. However, the combination of genome instability and high proliferation rates cause mitotic stress, which selects for tumor cells that can adapt and survive under these conditions, because even tumor cells have a loss of viability if aneuploidy exceeds a certain threshold ^48, 49^. One mechanism by which tumor cells adapt to prevent mitotic catastrophe is by upregulating SAC genes to ensure that mitosis is not completed until chromosomes are fully attached to the microtubule spindles and can segregate properly ^50^. Tumor cells with high SAC function are more resistant to drugs that cause DNA damage and mitotic stress because these cells are more likely to arrest and repair the damage before dividing^51^. The expression of SAC genes is frequently elevated in TNBC, the model used in our study, and high expression is associated with poor overall outcomes ^52–54^. TNBC is characterized by high chromosome instability and proliferation rate and requires an active SAC to maintain viability ^12–14^. The fact that IRS2 expression is elevated in TNBC and that it contributes to tumor cell viability in response to mitotic stress identifies a potential IRS2-specific vulnerability that could be exploited therapeutically in this breast cancer subtype.

## Method and Materials

### Cells, antibodies, and reagents

SUM-159 cells that were authenticated by STR profiling at the University of Arizona Genetics Core in February 2022 were a kind gift from Art Mercurio (UMass Chan Medical School, Worcester, MA) and were grown in F12 media (Gibco) containing 5% FBS (Sigma-Aldrich), 5 µg/mL insulin (Sigma-Aldrich) and 1 µg/mL hydrocortisone (Sigma-Aldrich). MDA-MB-231 cells were obtained from the ATCC Cell Biology Collection and were grown in RPMI (Gibco) containing 10% FBS (Sigma). All cells were screened regularly and tested negative for mycoplasma by PCR (Abm).

IRS2 (*IRS2^-/-^*) knockout cells were generated by CRISPR/Cas9-mediated gene editing by electroporation of a ribonucleoprotein (RNP) complex of CRISPR-gRNA and Cas9 protein prepared according to manufacturer’s protocol (IDT). Guide RNAs used were as follows: sgIRS2-1, GAG AAC TGC ACG GTG ATC GA; sgIRS2-2, TCG GTA ATG ACC GTG AGC TT. The control non-targeting guide RNA (sgNT) was purchased from IDT.

Human IRS2 was a kind gift from Adrian Lee (University of Pittsburgh, Pittsburgh, PA). cDNA was subcloned into the pCDH-CMV-MCS-EF1-puro lentiviral vector (System Bioscience) with a C-terminal FLAG tag ^7^. S731A and S736A mutations were introduced into the pCDH-hIRS2-FFH-EF1 plasmid by site directed mutagenesis using the primers 5’–GAG AGC GCC CCC GAG GAC AGT GGG TAC ATG -3’ and 5’–GGC GGG CGC GCT GGC CTT GTA GCC GCC -3’ according to the manufacturer’s protocol (Q5 Site Directed Mutagenesis Kit; New England Biolabs). The IRS2-Y5F mutant construct was generated by Genewiz and subcloned into the pCDH-CMV-MCS-EF1-puro lentiviral vector (System Bioscience) with a C-terminal FLAG tag. Lentiviruses were generated by co-transfection of IRS2 plasmids and packaging plasmids (pMD2.G and psPAX2) into HEK293FT cells using lipofectamine 3000 according to the manufacturer’s instruction. Cells were infected for 24 h with virus in the presence of 8 ug/ml polybrene (Sigma-Aldrich) and selected with 2 µg/mL puromycin (Gold Biotechnology). pLenti6-H2B-mCherry was a gift from Torsten Wittmann (Addgene plasmid # 89766; http://n2t.net/addgene:89766; RRID:Addgene_89766) ^25^. Lentivirus was generated as described above and infected cells were selected with 10 ug/ml blasticidin (Invivogen).

Primary antibodies used in this study include: rabbit anti-IRS2 (Western blotting [WB], 1:1000; Cell Signaling Technology [CST]; # 4502; RRID:AB_2125774), rabbit anti-IRS2 (Immunofluorescence [IF], 1:200; Abcam; # ab134101), Alexa Fluor594 conjugated anti-IRS2 (Flow cytometry, 1:50; Santa Cruz, #sc390761), rabbit anti-PLK1 (WB, 1:1000; CST; #4513), mouse anti-PLK1 (IF, 1:600; Immunoprecipitation [IP], 1:800; Invitrogen; #37-7000), rabbit anti-pPLK1 (T210) (WB, 1:1000; CST; #5472), rabbit anti-γ tubulin (IF, 1:200; Abclonal; #A9657), human anti-CREST (IF, 1:200; Antibodies Incorporated; #15-234), rabbit anti-BUBR1 (WB, 1:2000; IP, 1:1000; Abclonal; #14525), mouse anti-CYCLIN A (WB, 1:2000; Santa Cruz; #271682), rabbit anti-CYCLIN B1 (WB, 1:2000; CST; #4138), mouse anti-EMI1 (WB, 1:1000; Invitrogen; #37-6600), mouse anti-ACTIN (WB, 1:2000; Thermo Fisher Scientific; # MA5-11869; RRID:AB_11004139), mouse anti-GAPDH (WB, 1:2000; Santa Cruz Biotechnology; # sc-32233; RRID:AB_627679), mouse anti-Flag (WB, 1:2000; Sigma-Aldrich; # F3165; RRID:AB_259529), Alexa Fluor488 conjugated anti phospho histone H3 (1:200; Biolegend, #650803).

### Cell cycle synchronization

For single thymidine block, release and nocodazole treatment, cells were treated with thymidine (2 mM, Sigma-Aldrich) for 20 hrs, released into normal culture media for 8 hrs, and then treated with nocodazole (90 ng/ml, Sigma-Aldrich) for 16 hrs. Mitotic cells were collected by shake-off and re-plated in growth media for 5 hrs. To arrest cells in G_1_, cells were treated with 500nM (SUM-159 cells) or 150nM (MDA-MB-231 cells) Palbociclib for 24 hrs. To arrest cells at the G_1_/S boundary, cells were treated with 2 mM thymidine for 20 hrs, released into normal culture media for 8 hrs, and then subjected to a second thymidine treatment for 16 hrs. To arrest cells at the G_2_/M boundary, cells were treated with Ro3306 (10 μM, Selleckchem) for 20 hrs.

### Immunoprecipitation and immunoblotting

For whole-cell extract immunoblots, cells were solubilized at 4 °C in lysis buffer (20 mM Tris buffer, pH 7.4, 1% Nonidet P-40, 0.137 M NaCl, 10% glycerol) containing phosphatase inhibitors (Roche) and protease inhibitors (Roche). For treatment with λ phosphatase, cells were lysed in buffer without phosphatase inhibitors and cell extracts were treated with λ phosphatase (Santa Cruz) for 1 hr at 30 °C. For immunoprecipitations, cells were extracted in lysis buffer (50 mM Tris buffer, pH 7.4, 1% Triton X-100, 150 mM NaCl) containing phosphatase inhibitors (Roche) and protease inhibitors (Roche). Aliquots of cell extracts containing equivalent amounts of protein were precleared for 1 hr with IgG and protein G-Sepharose beads (GE Healthcare) or protein A agarose beads (Thermo) and then incubated for 3 hrs or overnight at 4 °C with specific antibodies and beads with constant agitation. For IP with FLAG antibody, anti-FLAG M2 affinity gel (Sigma-Aldrich; #A2220) was used. The beads were washed three times in extraction buffer.

Cell extracts containing equivalent amounts of protein or immune complexes were resolved by SDS-PAGE and transferred to nitrocellulose membranes. Membranes were blocked using 5% non-fat milk in 1x TBST (50 mM Tris, pH 7.5, 150 mM NaCl, 0.01% Tween 20). Primary and secondary antibody incubations were carried out in 3% bovine serum albumin (BSA) or 5% non-fat milk in 1X TBST overnight at 4 °C or 1 h at room temperature, respectively. Bands were detected by chemiluminescence using a ChemiDoc XRS+ system (Bio-Rad). For immunoblot quantification, the signals from the protein of interest were normalized with that of the loading control using Image J.

### Quantitative polymerase chain reaction (qPCR)

RNA was extracted using the RNA-easy kit (Qiagen). cDNA was synthesized using an AzuraQuant cDNA synthesis kit (Azura Genomics). qPCR was performed in an Applied Biosystems QuantStudio 6 Flex apparatus using AzuraView GreenFast qPCR Blue Mix (Azura Genomics). The delta –delta Ct method was used to determine relative mRNA expression. Primers used were as follows: h*IRS2*-Fwd 5’-CCA CAT CGT GAA AGA GTG A-3’; h*IRS2*-Rev 5’-CAG AGT CCA CAG ATG TTT CCA A-3’; r18s-Fwd 5’-GTC GCT CGC TCC TCT CCT ACT-3’; r18s-Rev 5’-TCT GAT AAA TGC ACG CAT CCC-3’.

### Flow cytometry

For cell cycle analysis, adherent cells were collected by trypsinization and combined with non-adherent cells from the culture medium. Cells were washed with PBS then fixed in ice-cold 70% ethanol overnight at 4 °C. The fixed cells were washed with PBS twice before treatment with 1 ug/ml DAPI in PBS for 20 min at room temperature. To detect IRS2 or phospho histone H3, fixed cells were permeabilized by 0.2% tween-20 in PBS treatment for 10 min before incubation with antibodies for 30 min at room temperature. The cells were analyzed by flow cytometry using a ZE5 cell analyzer (Bio-Rad). The data was processed and analyzed using FlowJo software (v.10.10).

### Mitotic/nuclear fragmentation analysis

Cells were plated in 12 or 24-well plates and treated with nocodazole (90 ng/ml). 10 or 30 hrs after the treatment, live cells were immediately imaged using an EVOS cell imaging system (Thermo Fisher). Mitotic cells or cells with fragmented nuclei were counted based on nuclear imaging pattern and 200-400 cells per well were analyzed.

### Time-lapse imaging

H2B-mCherry expressing cells were plated in a 96 well plate and incubated overnight. Cells were treated with compounds and imaged immediately afterward. All images were collected using an Incucyte imaging system (Sartorius). Three fields of view were collected per well, and phase contrast and fluorescence images were captured at 10 min intervals for 24 hr.

### Immunofluorescence

Cells for immunofluorescence imaging were plated in 8 well chamber slides (Ibidi) or coverslips placed in 24-well plates. Cells were fixed with 4% PFA for 10 min and permeabilized with 0.1% Triton X-100 for 10 min followed by blocking with 1% BSA, 0.3% Triton X-100 in PBS for 30 min at RT. Cells were incubated with primary antibodies (diluted in blocking buffer) for 2-3 hours at RT followed by washing with PBS and incubated with AlexaFluor conjugated secondary antibodies in the dark for 1h with Hoechst 33342 (2 ug/ml). Following washing steps, PBS was added to the chambers and images were collected using a Nikon A1 confocal microscope. Quantification of the signal intensity for the line scan analysis was achieved using Image J. For experiments using DFN-IRS2-SUM159 cells, cells were fixed with 100% methanol at -20 °C for 5 minutes and processed as described above.

### LC-MS spectrometry

IRS1^-/-^,IRS2^-/-^ SUM159 cells expressing FLAG-tagged IRS2-WT were serum starved and then stimulated with insulin (500 ng/ml) for 15 min. Cells were extracted in lysis buffer (50 mM Tris, pH 7.4, 150 mM NaCl, 1 mM EDTA, 1% Triton X-100, containing phosphatase inhibitors (Roche) and protease inhibitors (Roche) and cell extracts were clarified by centrifugation (13000 rpm for 15 min at 4 °C). Cell extracts (2mg) were incubated with agarose-conjugated anti-FLAG antibody (Sigma-Aldrich, 45 ul per IP) for 3 hrs at 4 °C, then washed three times with lysis buffer. Samples were then incubated with 3x FLAG peptide (0.25 mg/ml, 50 ul per IP, 30 min at 4 °C) to elute the immunoprecipitated proteins. The elution step was performed twice for each sample and then eluents were combined to a single tube.

Proteins from pull-downs were precipitated using 20% tri-chloroacetic acid (TCA), washed twice with acetone (Burdick & Jackson, Muskegon, MI), and digested overnight in 25 mM ammonium bicarbonate with trypsin (Promega) for mass spectrometric analysis. Peptides were analyzed on a Q-Exactive Plus quadrupole equipped with Easy-nLC 1000 (ThermoScientific) and nanospray source (ThermoScientific). Peptides were resuspended in 5% methanol / 1% formic acid and analyzed as previously described ^55^. Raw data were searched using COMET (release version 2014.01) in high resolution mode ^56^ against a target-decoy (reversed) ^57^ version of the human proteome sequence database (UniProt; 40704 entries of forward and reverse protein sequences) with a precursor mass tolerance of +/- 1 Da and a fragment ion mass tolerance of 0.02 Da, and requiring fully tryptic peptides (K, R; not preceding P) with up to three mis-cleavages. Searches were filtered using orthogonal measures including mass measurement accuracy (+/- 3 ppm), Xcorr for charges from +2 through +4, and dCn targeting a <1% FDR at the peptide level. Quantification of LC-MS/MS spectra was performed using MassChroQ ^58^ and the iBAQ method^59^. Mass spectrometry data have been deposited to ProteomeXchange: PXD052524 and MSV000094848: https://massive.ucsd.edu/ProteoSAFe/dataset.jsp?task=407565b7b7674af886f18cdd1aaef809. password: 67631

### Statistical analysis

Statistical analysis was performed using Prism7, Graphpad. Student’s t-test (two-sided) was applied and p-value of <0.05 was considered to indicate statistical significance. For comparison of more than 2 groups, ordinary one-way ANOVA followed by Tukey’s multiple comparisons was applied.

## Acknowledgements

This work was supported by National Institute of Health (NIH) R01 grants CA229910, CA227993 and CA290778 (LMS) and R35 grant GM119455 (ANK). The content is solely the responsibility of the authors and does not necessarily represent the official views of the National Institutes of Health.

## Author Contributions

J-SL and LMS were involved in the conception and design of the project and wrote the manuscript; J-SL, QTB, MJ, JSM, IN and ANK were involved with the development of methodology and the acquisition and analysis of data. All authors reviewed and approved the final version of the manuscript.

