## Supplemental Files for "Interaction of IRS2 with PLK1 protects cells from mitotic stress"

### Figure S1

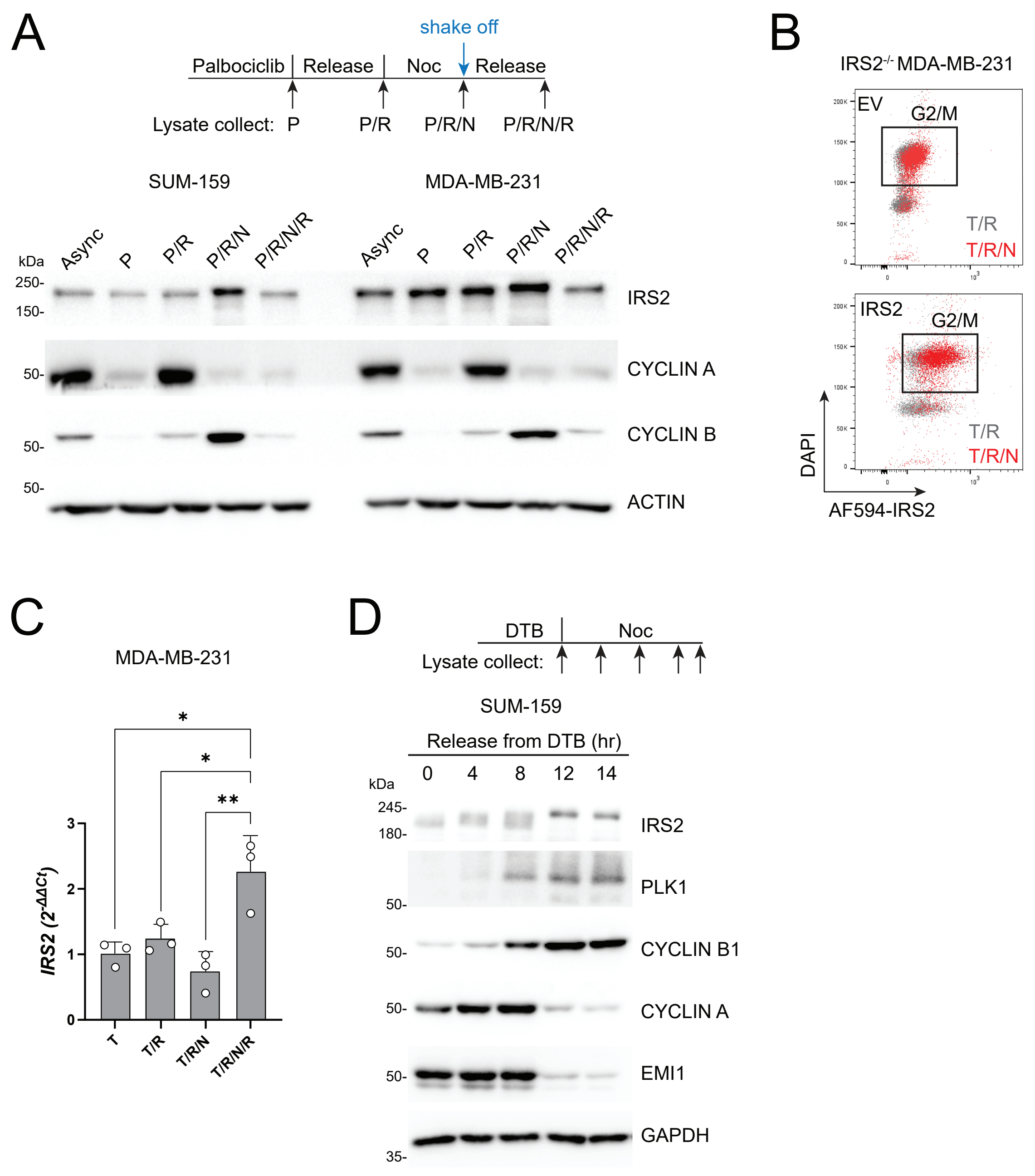

A
shake off
Palbociclib
Release
Noc
Release
Lysate collect: P
P/R
P/R/N
P/R/N/R
SUM-159
MDA-MB-231
kDa
Async
P
P/R
P/R/N
P/R/N/R
250-
150-
IRS2
50-
CYCLIN A
50-
CYCLIN B
50-
ACTIN
B
IRS2-/- MDA-MB-231
EV
G2/M
T/R
T/R/N
IRS2
G2/M
T/R
T/R/N
DAPI
AF594-IRS2
C
MDA-MB-231
IRS2 (2-ΔΔCt)
T
T/R
T/R/N
T/R/N/R
D
DTB
Noc
Lysate collect:
SUM-159
Release from DTB (hr)
0
4
8
12
14
kDa
245-
180-
IRS2
PLK1
50-
50-
CYCLIN B1
50-
CYCLIN A
50-
EMI1
GAPDH
35-

### Figure S2

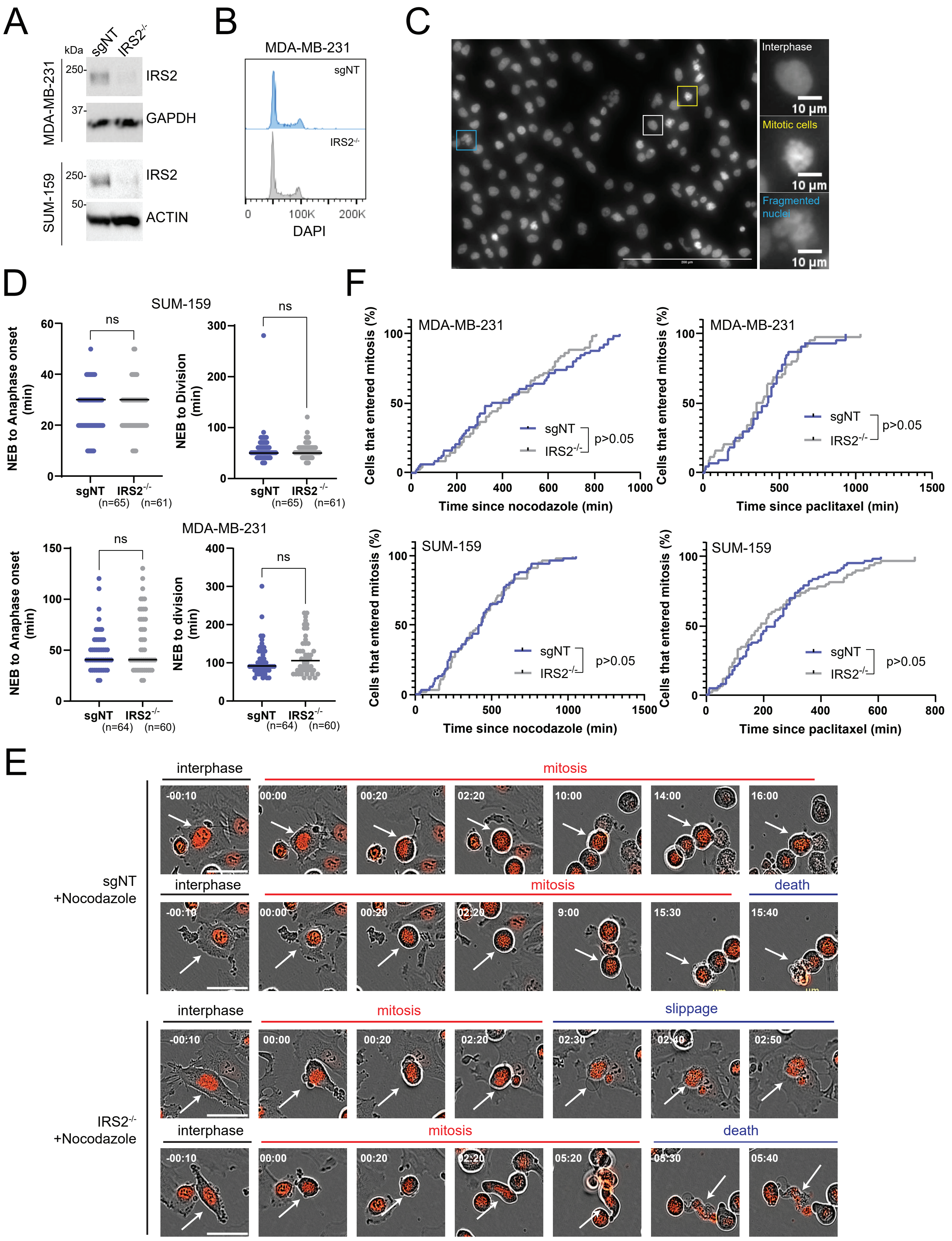
A
kDa
sgNT
IRS2-/-
MDA-MB-231
250-
37-
IRS2
GAPDH
SUM-159
250-
50-
IRS2
ACTIN
B
MDA-MB-231
sgNT
IRS2-/-
0
100K
200K
DAPI
C
Interphase
10 µm
Mitotic cells
10 µm
Fragmented nuclei
10 µm
D
SUM-159
ns
NEB to Anaphase onset (min)
sgNT
IRS2-/-
(n=65)
(n=61)
NEB to Division (min)
MDA-MB-231
NEB to division (min)
(n=64)
(n=60)
F
MDA-MB-231
Cells that entered mitosis (%)
Time since nocodazole (min)
Time since paclitaxel (min)
p>0.05
SUM-159
E
interphase
mitosis
death
slippage
sgNT +Nocodazole
IRS2-/- +Nocodazole

### Figure S3

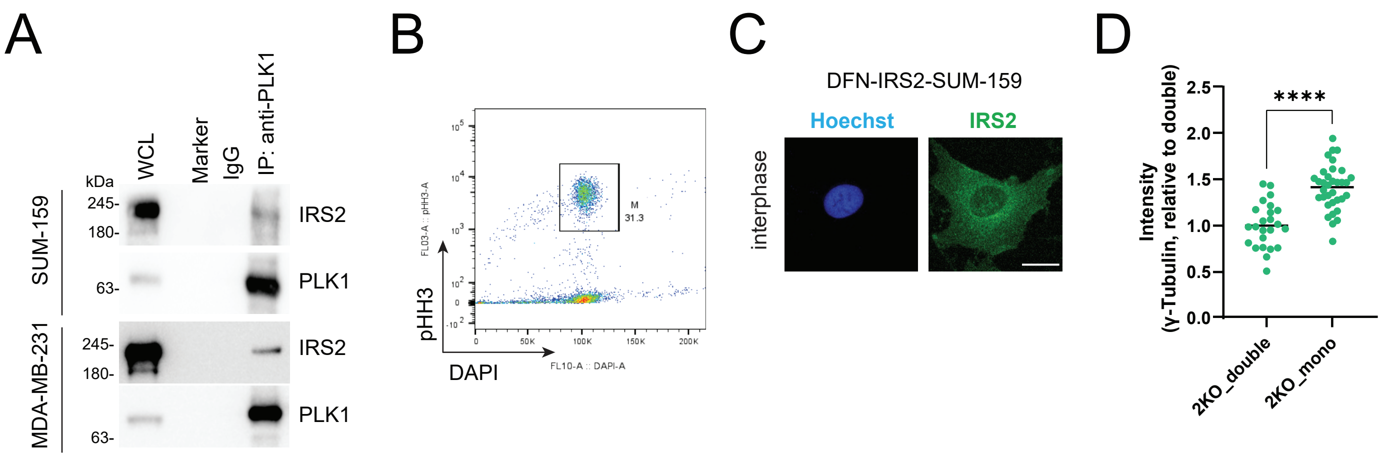
A
WCL
Marker
IgG
IP: anti-PLK1
kDa
245-
180-
63-
SUM-159
MDA-MB-231
IRS2
PLK1
B
pHH3
DAPI
M
31.3
C
DFN-IRS2-SUM-159
Hoechst
IRS2
interphase
D
Intensity
(γ-Tubulin, relative to double)
****
2KO_double
2KO_mono

### Figure S4

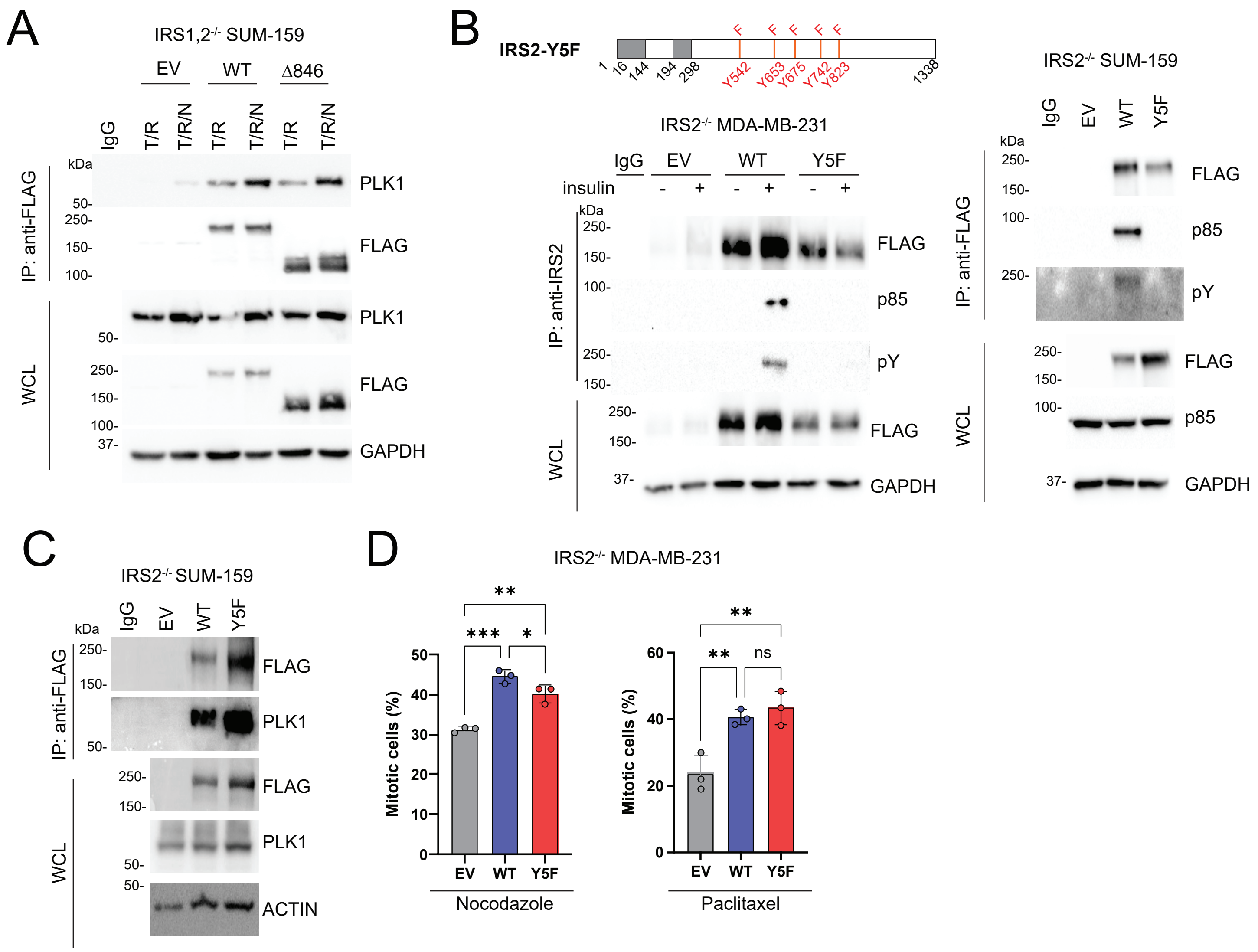

A
IRS1,2-/- SUM-159
EV
WT
Δ846
IgG
T/R
T/R/N
kDa
IP: anti-FLAG
WCL
PLK1
FLAG
GAPDH
B
IRS2-Y5F
Y542
Y653
Y675
Y742
Y823
1338
IRS2-/- MDA-MB-231
insulin
IP: anti-IRS2
p85
pY
IRS2-/- SUM-159
Y5F
C
ACTIN
D
IRS2-/- MDA-MB-231
Mitotic cells (%)
Nocodazole
Paclitaxel
ns

### Figure S5

A

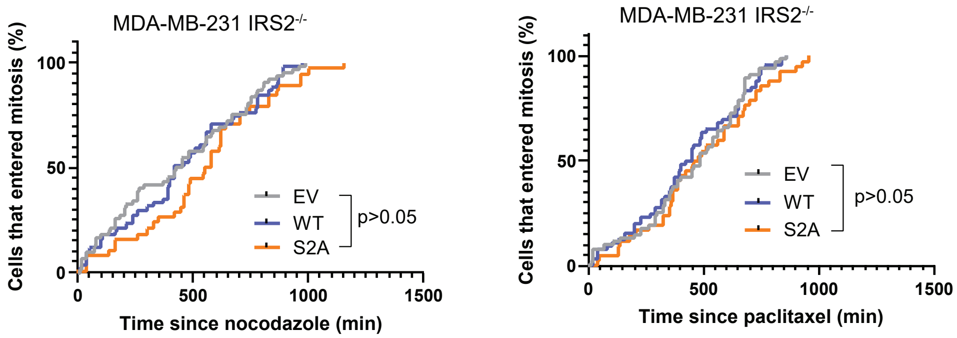
MDA-MB-231 IRS2-/-
Cells that entered mitosis (%)
100
50
0
0
500
1000
1500
Time since nocodazole (min)
EV
WT
S2A
p>0.05
MDA-MB-231 IRS2-/-
Cells that entered mitosis (%)
100
50
0
0
500
1000
1500
Time since paclitaxel (min)
EV
WT
S2A
p>0.05

#### Supplementary Figure Legends

**Figure S1.** (A) SUM-159 (500 nM) and MDA-MB-231 (150 nM) cells were treated with palbociclib for 24 hrs (P), released for 8 hrs (P/R) then treated with nocodazole (90 ng/ml) for 16 hrs. Mitotic cells were collected by shake-off (P/R/N) then replated and released into growth media for 5 hrs (P/R/N/R). Cell extracts were analyzed by immunoblotting for IRS2 and cell cycle proteins. (B) *IRS2*$^{-/-}$ MDA-MB-231 cells expressing empty vector (EV) or IRS2-WT were treated as shown in Fig 1A. T/R or T/R/N cells were collected and fixed, and IRS2 expression was analyzed by flow cytometry. (C) MDA-MB-231 cells were treated with thymidine (2 mM) for 18 hrs (T), released for 8 hrs (T/R) then treated with nocodazole (90 ng/ml) for 16 hrs. Mitotic cells were collected by shake-off (T/R/N) then replated and released into growth media for 5 hrs (T/R/N/R). The data shown represent the mean ± S.D. of *IRS2* mRNA expression from three independent experiments. (D) SUM-159 cells were synchronized by double thymidine block and released into growth media in the presence of nocodazole (90 ng/ml). Cell extracts were analyzed by immunoblotting for IRS2 and cell cycle proteins.

**Figure S2.** (A) MDA-MB-231 or SUM-159 cells treated with non-targeting guide RNA (*sgNT*) or IRS2 knockout (KO) cells (*IRS2*$^{-/-}$) were analyzed for IRS2 expression by immunoblotting. (B) Asynchronously grown cells were stained with DAPI and analyzed by flow cytometry for cell cycle profiles. (C) Representative images of H2B-mCherry labeled *IRS2*$^{-/-}$ MDA-MB-231 cells treated with nocodazole (90 ng/ml, 10 hrs) from Figures 2A and 3A showing interphase, mitotic and fragmented nuclei. (D) *sgNT* or *IRS2*$^{-/-}$ cells growing asynchronously in complete growth media were imaged every 10 minutes by widefield time-lapse microscopy for 24 hrs. Each dot represents the mitotic duration of an individual cell measured as the time from nuclear envelope breakdown to anaphase onset or completion of division. *sgNT* SUM-159, n=65; *IRS2*$^{-/-}$ SUM-159, n=61; *sgNT* MDA-MB-231, n=64; *IRS2*$^{-/-}$ MDA-MB-231, n=60. (E) Representative images of SUM-159 cells expressing H2B-mCherry from Figure 2B. Scale bar, 50 μm. Time=hr:min. (F) Quantification of time to mitosis-entry, with time from drug treatment to nuclear envelope breakdown plotted as cumulative frequency (from Figure 2B).

**Figure S3.** (A) SUM-159 and MDA-MB-231 cells were treated with nocodazole (90 ng/ml) for 16 hrs and cell extracts from mitotic cells were immunoprecipitated with PLK1-specific antibodies. (B) SUM-159 cells were arrested at the $G_2$/M boundary by Ro3306 (10 uM) treatment for 20 hrs then released into mitosis for 30 min. Cells were fixed and processed for analysis of phosphohistone-H3 by flow cytometry to confirm mitotic cells. (C) DFN-IRS2 SUM-159 cells were imaged for IRS2. Scale bar, 20 µm. (D) Quantification of γ-tubulin immunostaining intensity in $IRS2^{-/-}$ cells from the immunofluorescence images in Figure 3G.

**Figure S4.** (A) $IRS1,2^{-/-}$ SUM-159 cells expressing, EV, IRS2WT, IRS2Δ846 were synchronized as shown in Figure 1A. Cell extracts were immunoprecipitated with FLAG-specific antibodies and immunoblotted as labeled. (B) Cell extracts from cells stimulated with insulin (100 ng/ml, 10 min, left) or growing asynchronously in complete growth media (right) were immunoprecipitated with FLAG or IRS2-specific antibodies and immunoblotted as labeled. (C) $IRS2^{-/-}$ SUM-159 cells were treated with nocodazole (90 ng/ml) for 16 hrs and cell extracts from mitotic cells were immunoprecipitated with FLAG-specific antibodies. (D) *IRS2*$^{-/-}$ MDA-MB-231 cells expressing EV, IRS2-WT or IRS2-Y5F were treated with nocodazole (90 ng/ml) or paclitaxel (100 nM). Cells were imaged after 10 hours of treatment. The data shown represent the mean ± SD of a representative experiment performed three times independently.

**Figure S5.** (A) H2B-mCherry labeled *IRS2*$^{-/-}$ MDA-MB-231 cells expressing EV, IRS2-WT or IRS2-S2A were treated with nocodazole (90 ng/ml) or paclitaxel (100 nM). Cells were imaged every 10 minutes by widefield time-lapse microscopy for 24 hrs. Time to mitosis-entry was quantified, with time from drug treatment to nuclear envelope breakdown plotted as cumulative frequency (From Figure 4B).

| Table S1. Proteins identified and quantified in proteomics (Related to Fig 4A) | | | | |
|---|---|---|---|---|
| name | accession | description | Fold change (stim/unstim) | p-value |
| EIF3A_HUMAN | Q14152 | Eukaryotic translation initiation factor 3 subunit A OS=Homo sapiens OX=9606 GN=EIF3A PE=1 SV=1 | 4.43 | 0.0262 |
| RBM6_HUMAN | P78332 | RNA-binding protein 6 OS=Homo sapiens OX=9606 GN=RBM6 PE=1 SV=5 | 4.01 | 0.0025 |
| BAG3_HUMAN | O95817 | BAG family molecular chaperone regulator 3 OS=Homo sapiens OX=9606 GN=BAG3 PE=1 SV=3 | 3.76 | 0.0110 |
| CKAP5_HUMAN | Q14008 | Cytoskeleton-associated protein 5 OS=Homo sapiens OX=9606 GN=CKAP5 PE=1 SV=3 | 3.75 | 0.0056 |
| PRPS1_HUMAN | P60891 | Ribose-phosphate pyrophosphokinase 1 OS=Homo sapiens OX=9606 GN=PRPS1 PE=1 SV=2 | 3.16 | 0.0363 |
| SR140_HUMAN | O15042 | U2 snRNP-associated SURP motif-containing protein OS=Homo sapiens OX=9606 GN=U2SURP PE=1 SV=2 | 3.07 | 0.0116 |
| E2F7_HUMAN | Q96AV8 | Transcription factor E2F7 OS=Homo sapiens OX=9606 GN=E2F7 PE=1 SV=3 | 3.04 | 0.0045 |
| SF3B1_HUMAN | O75533 | Splicing factor 3B subunit 1 OS=Homo sapiens OX=9606 GN=SF3B1 PE=1 SV=3 | 2.99 | 0.0162 |
| NSRP1_HUMAN | Q9H0G5 | Nuclear speckle splicing regulatory protein 1 OS=Homo sapiens OX=9606 GN=NSRP1 PE=1 SV=1 | 2.98 | 0.0054 |
| SND1_HUMAN | Q7KZF4 | Staphylococcal nuclease domain-containing protein 1 OS=Homo sapiens OX=9606 GN=SND1 PE=1 SV=1 | 2.78 | 0.0271 |
| PLK1_HUMAN | P53350 | Serine/threonine-protein kinase PLK1 OS=Homo sapiens OX=9606 GN=PLK1 PE=1 SV=1 | 2.75 | 0.0331 |
| HNRPL_HUMAN | P14866 | Heterogeneous nuclear ribonucleoprotein L OS=Homo sapiens OX=9606 GN=HNRNPL PE=1 SV=2 | 2.67 | 0.0302 |
| ANM1_HUMAN | Q99873 | Protein arginine N-methyltransferase 1 OS=Homo sapiens OX=9606 GN=PRMT1 PE=1 SV=3 | 2.67 | 0.0163 |
| TET2_HUMAN | Q6N021 | Methylcytosine dioxygenase TET2 OS=Homo sapiens OX=9606 GN=TET2 PE=1 SV=3 | 2.51 | 0.0131 |
| POTEI_HUMAN | P0CG38 | POTE ankyrin domain family member I OS=Homo sapiens OX=9606 GN=POTEI PE=3 SV=1 | 2.31 | 0.0147 |
| ANFY1_HUMAN | Q9P2R3 | Rabankyrin-5 OS=Homo sapiens OX=9606 GN=ANKFY1 PE=1 SV=2 | 2.29 | 0.0131 |
| CHERP_HUMAN | Q8IWX8 | Calcium homeostasis endoplasmic reticulum protein OS=Homo sapiens OX=9606 GN=CHERP PE=1 SV=3 | 2.29 | 0.0485 |
| P85A_HUMAN | P27986 | Phosphatidylinositol 3-kinase regulatory subunit alpha OS=Homo sapiens OX=9606 GN=PIK3R1 PE=1 SV=2 | 2.28 | 0.0042 |
| PABP4_HUMAN | Q13310 | Polyadenylate-binding protein 4 OS=Homo sapiens OX=9606 GN=PABPC4 PE=1 SV=1 | 2.25 | 0.0092 |
| PLEC_HUMAN | Q15149 | Plectin OS=Homo sapiens OX=9606 GN=PLEC PE=1 SV=3 | 2.22 | 0.0467 |
| NONO_HUMAN | Q15233 | Non-POU domain-containing octamer-binding protein OS=Homo sapiens OX=9606 GN=NONO PE=1 SV=4 | 2.20 | 0.0092 |
| HS74L_HUMAN | O95757 | Heat shock 70 kDa protein 4L OS=Homo sapiens OX=9606 GN=HSPA4L PE=1 SV=3 | 2.15 | 0.0118 |
| AKAP2_HUMAN | Q9Y2D5 | A-kinase anchor protein 2 OS=Homo sapiens OX=9606 GN=AKAP2 PE=1 SV=3 | 2.14 | 0.0279 |
| SF3B2_HUMAN | Q13435 | Splicing factor 3B subunit 2 OS=Homo sapiens OX=9606 GN=SF3B2 PE=1 SV=2 | 2.12 | 0.0303 |
| POTEE_HUMAN | Q6S8J3 | POTE ankyrin domain family member E OS=Homo sapiens OX=9606 GN=POTEE PE=2 SV=3 | 2.12 | 0.0221 |
| POTEF_HUMAN | A5A3E0 | POTE ankyrin domain family member F OS=Homo sapiens OX=9606 GN=POTEF PE=1 SV=2 | 2.12 | 0.0221 |
| POTEJ_HUMAN | P0CG39 | POTE ankyrin domain family member J OS=Homo sapiens OX=9606 GN=POTEJ PE=3 SV=1 | 2.12 | 0.0080 |
| SPTN2_HUMAN | O15020 | Spectrin beta chain, non-erythrocytic 2 OS=Homo sapiens OX=9606 GN=SPTBN2 PE=1 SV=3 | 2.09 | 0.0374 |
| ACTC_HUMAN | P68032 | Actin, alpha cardiac muscle 1 OS=Homo sapiens OX=9606 GN=ACTC1 PE=1 SV=1 | 1.99 | 0.0194 |
| ACTS_HUMAN | P68133 | Actin, alpha skeletal muscle OS=Homo sapiens OX=9606 GN=ACTA1 PE=1 SV=1 | 1.99 | 0.0194 |
| ACTB_HUMAN | P60709 | Actin, cytoplasmic 1 OS=Homo sapiens OX=9606 GN=ACTB PE=1 SV=1 | 1.97 | 0.0341 |
| ACTG_HUMAN | P63261 | Actin, cytoplasmic 2 OS=Homo sapiens OX=9606 GN=ACTG1 PE=1 SV=1 | 1.97 | 0.0341 |
| JAK1_HUMAN | P23458 | Tyrosine-protein kinase JAK1 OS=Homo sapiens OX=9606 GN=JAK1 PE=1 SV=2 | 1.96 | 0.0196 |
| DHX15_HUMAN | O43143 | Pre-mRNA-splicing factor ATP-dependent RNA helicase DHX15 OS=Homo sapiens OX=9606 GN=DHX15 PE=1 | 1.96 | 0.0469 |
| HNRPK_HUMAN | P61978 | Heterogeneous nuclear ribonucleoprotein K OS=Homo sapiens OX=9606 GN=HNRNPK PE=1 SV=1 | 1.95 | 0.0308 |
| ACTA_HUMAN | P62736 | Actin, aortic smooth muscle OS=Homo sapiens OX=9606 GN=ACTA2 PE=1 SV=1 | 1.94 | 0.0304 |
| ACTH_HUMAN | P63267 | Actin, gamma-enteric smooth muscle OS=Homo sapiens OX=9606 GN=ACTG2 PE=1 SV=1 | 1.94 | 0.0304 |
| PABP3_HUMAN | Q9H361 | Polyadenylate-binding protein 3 OS=Homo sapiens OX=9606 GN=PABPC3 PE=1 SV=2 | 1.85 | 0.0346 |
| P85B_HUMAN | O00459 | Phosphatidylinositol 3-kinase regulatory subunit beta OS=Homo sapiens OX=9606 GN=PIK3R2 PE=1 SV=2 | 1.85 | 0.0493 |
| SRC8_HUMAN | Q14247 | Src substrate cortactin OS=Homo sapiens OX=9606 GN=CTTN PE=1 SV=2 | 1.58 | 0.0040 |
| F263_HUMAN | Q16875 | 6-phosphofructo-2-kinase/fructose-2,6-bisphosphatase 3 OS=Homo sapiens OX=9606 GN=PFKFB3 PE=1 SV=1 | 1.30 | 0.0399 |
| TR150_HUMAN | Q9Y2W1 | Thyroid hormone receptor-associated protein 3 OS=Homo sapiens OX=9606 GN=THRAP3 PE=1 SV=2 | 1.53 | 0.0122 |
